# An apicoplast localized GTPase is essential for *Toxoplasma gondii* survival

**DOI:** 10.1101/2025.09.11.675638

**Authors:** Michael B. Griffith, Morgan E. Wagner, Victoria L. Robinson, Aoife T. Heaslip

## Abstract

The apicoplast is an essential organelle found in Apicomplexa, a large phylum of intracellular eukaryotic pathogens. The apicoplast produces metabolites that are utilized for membrane biogenesis and energy production. A majority of apicoplast resident proteins are encoded by the nuclear genome and are trafficked to the apicoplast, and are referred to as <u>N</u>uclear <u>E</u>ncoded and <u>A</u>picoplast <u>T</u>argeted (NEAT) proteins. In this study, we characterized a NEAT protein named TgBipA, that is a homolog of the highly conserved prokaryotic translational GTPase BipA. BipA is essential for bacterial survival in stress conditions and functions through interactions with the prokaryotic ribosome, although its role is not fully understood. Through genetic knockouts of TgBipA and immunofluorescence imaging we show that loss of TgBipA results in apicoplast genome replication defects, disruption of NEAT trafficking, loss of the apicoplast, and subsequently parasite death. Furthermore, we show through comparative studies that this phenotype closely resembles the delayed death phenomenon observed when inhibiting apicoplast translation. Finally, we show that TgBipA is an active GTPase *in vitro,* and its GTP hydrolysis activity is critical for its cellular function. Our findings demonstrate TgBipA is a GTPase which has an essential role in apicoplast maintenance, providing new insight into the cellular processes of the organelle.

**Importance Statement:** *Toxoplasma gondii,* and many other parasites in the phylum Apicomplexa, are pathogens with significant medical and veterinary importance. Most Apicomplexa contain a non-photosynthetic plastid organelle named the apicoplast. This organelle produces essential metabolites and perturbation of apicoplast function results in parasite death. The apicoplast contains bacterial-like pathways for apicoplast genome replication and expression. Thus, the discovery of the apicoplast lead to optimism that this organelle would provide a wealth of anti-parasitic drug targets. Therefore, the identification and characterization of new apicoplast proteins could provide new opportunities for therapeutic development. In this study we characterized the function of a protein called TgBipA, a homolog of a highly conserved bacterial GTPase bipA which has been implicated in maturation of the 50S ribosomal subunit and adaptation to cellular stress. We show that TgBipA is essential for apicoplast maintenance and parasite survival.

## Introduction

The Apicomplexan phylum is a diverse group of obligate intracellular parasites including *Toxoplasma gondii*, *Plasmodium* spp., and *Babesia spp.*; the causative agents of human diseases toxoplasmosis, malaria, and babesiosis respectively^1–4^. *T. gondii* can cause life threatening disease when infection occurs *in utero* or in the immunocompromised^1,2,5,6^. For a successful infection, the parasite must survive within cells of its host where it relies on nutrients acquired from the host cell, and metabolites formed *de novo* ^7–9^. *T. gondii*, along with many other Apicomplexans, rely on a non-photosynthetic plastid organelle, termed the apicoplast, to produce essential metabolites^9^.

The apicoplast is a four membraned organelle derived from secondary endosymbiosis of a red algae, and thus is prokaryotic in origin^9^. The apicoplast houses four essential metabolic pathways; a fatty acid type II (FASII)^10^, heme precursor^11^, Isopentyl pyrophosphate (IPP)^12^, and lysophosphatidic acid (LPA) pathways^13^. In addition iron-sulfur cluster biosynthesis and ferredoxin redox pathways provide cofactors for metabolic enzymes^12,14–16^.

Most apicoplast proteins are nuclear encoded, the so-called <u>n</u>uclear <u>e</u>ncoded <u>a</u>picoplast trafficked (NEAT) pool of proteins^17–19^. Thus, organelle function requires trafficking these proteins to the apicoplast. Many NEAT proteins contain an N-terminal bipartite signaling motif which consists of a signal peptide (SP) which direct the protein to the ER during translation and an apicoplast transit peptide^17,20^. Removal of the SP in the ER reveals the apicoplast targeting sequence^18^. The apicoplast transit peptide can vary greatly in length and sequence, but tends to have an overall positive charge^17,20–22^. This variation has led to difficulties in identifying putative transit peptides through the use of prediction algorithms^23,24^. Other NEAT proteins contain neither an SP nor an identifiable transit peptide. What sequences guide these proteins to the ER, and subsequently the apicoplast is unclear^21,25–29^. Membrane associated, transmembrane domain (TM) containing NEAT proteins traffic to the apicoplast through ER-derived vesicular transport that occurs in a cell cycle dependent manner^30,31^. How lumen proteins are trafficked is poorly understood as these proteins are not observed in trafficking vesicles in wildtype parasites. Once NEAT lumen proteins have traversed the outermost apicoplast membrane, proteins are then trafficked across the three inner most membranes using a modified ERAD complex and TIC and TOC protein translocons^28,29,32^. Translocation into the lumen is followed by cleavage of the transit peptide to form the fully mature protein^33,34^.

In addition to the nuclear-encoded proteome, the apicoplast contains its own 35Kb genome^35^ encoding tRNAs, some 70S ribosomal components, RNA polymerase components, and two genes not directly involved in translation; *SufB* part of the an iron-sulfur cluster (ISC) pathway^14^ and *ClpC* which is speculated to be involved in protein translocation into the apicoplast lumen^35–37^. Despite the low number of metabolic proteins translated from the apicoplast genome, several antibiotics that target bacterial translation also have anti-parasitic effects in *Toxoplasma* and other Apicomplexa and are in clinical use, highlighting the importance of apicoplast translation^38–42^. Unfortunately, the utility of apicoplast translation targeting drugs has been somewhat limited by the so called delayed death effect whereby there are no apparent growth defects in the first intracellular growth cycle, but parasite death begins in the second intracellular cycle^42–45^. Although the mechanism of action of these antibiotics is well characterized in bacteria, it is poorly understood in parasites.

We recently identified a homolog of the bacterial GTPase BipA, TgBipA (TgME49_301380). It is 65% similar to its bacterial counterpart (bBipA). Although the exact function of bBipA is not understood it has been implicated in maturation of the 50S ribosomal subunit as well as other adaptive processes associated with stress ^46,47,51–55^. To study TgBipA, an inducible knockout parasite line was created. We show that loss of TgBipA and apicoplast-translation inhibition by doxycycline exhibit a similar cascade of phenotypic effects starting with a disruption to apicoplast genome maintenance. Next, vesicles containing NEAT proteins accumulate adjacent to the apicoplast. This coincides with a decrease in apicoplast size and eventual loss of the organelle. While parasite death begins in the ∼72 hours after translation inhibition or loss of TgBipA, we were surprised to observe parasites that survived for up to 9 days demonstrating that parasite death does not occur strictly in the second lytic cycle, as previously reported for the delayed death effect^37,43,48–50^. Finally, we demonstrate that TgBipA is an active GTPase and that its ability to hydrolyze GTP is critical for its function. Collectively, these results show that TgBipA plays an essential role in apicoplast maintenance.

## Results

### TgME49_301380 is an apicoplast localized homolog of bBipA

To define the biochemical and structural features of TgBipA we performed sequence alignments between *Salmonella typhimurium* bBipA and TgME49_301380 (Fig. 1A, Fig S1)^51^. TgBipA differs from bBipA in that it contains three insertions, two are serine rich (residues 215-256, 308-394) and the other is arginine rich (residues 642-714). It also has a ∼520 amino acid N-terminal extension that likely contains the signaling motifs required for apicoplast trafficking^17^, although the protein does not contain an identifiable signaling peptide based on primary sequence. TgBipA (amino acids 521-1333), shares 41% identity with bBipA^52^. The four conserved G-motifs are readily recognizable within the GTPase (G-) domain of TgBipA which shares 55% identity with the G-domain of bBipA ^53^. To gain insight into the structural features of the protein, we modeled TgBipA using Alphafold (AF) (Fig. 1B)^54^. The predicted structure suggests that the core of the protein, defined by five domains including a G-domain, OB fold, two alpha beta domains and a C-terminal domain unique to the BipA family, are similar in both proteins. The pLDDT scores, which denote the confidence of the prediction, are high throughout this part of the model (> 0.9). In contrast, AF was unable to define any structural features in the three large insertions (pLDDT < 50). This low confidence score suggested they are intrinsically disordered (IDR), subsequently confirmed by a PONDR-FIT^55^ assessment of the protein (Fig. S1). The inherent flexibility of these regions likely provides multiple conformational states important to the cellular functioning of TgBipA.

**Fig. 1.**
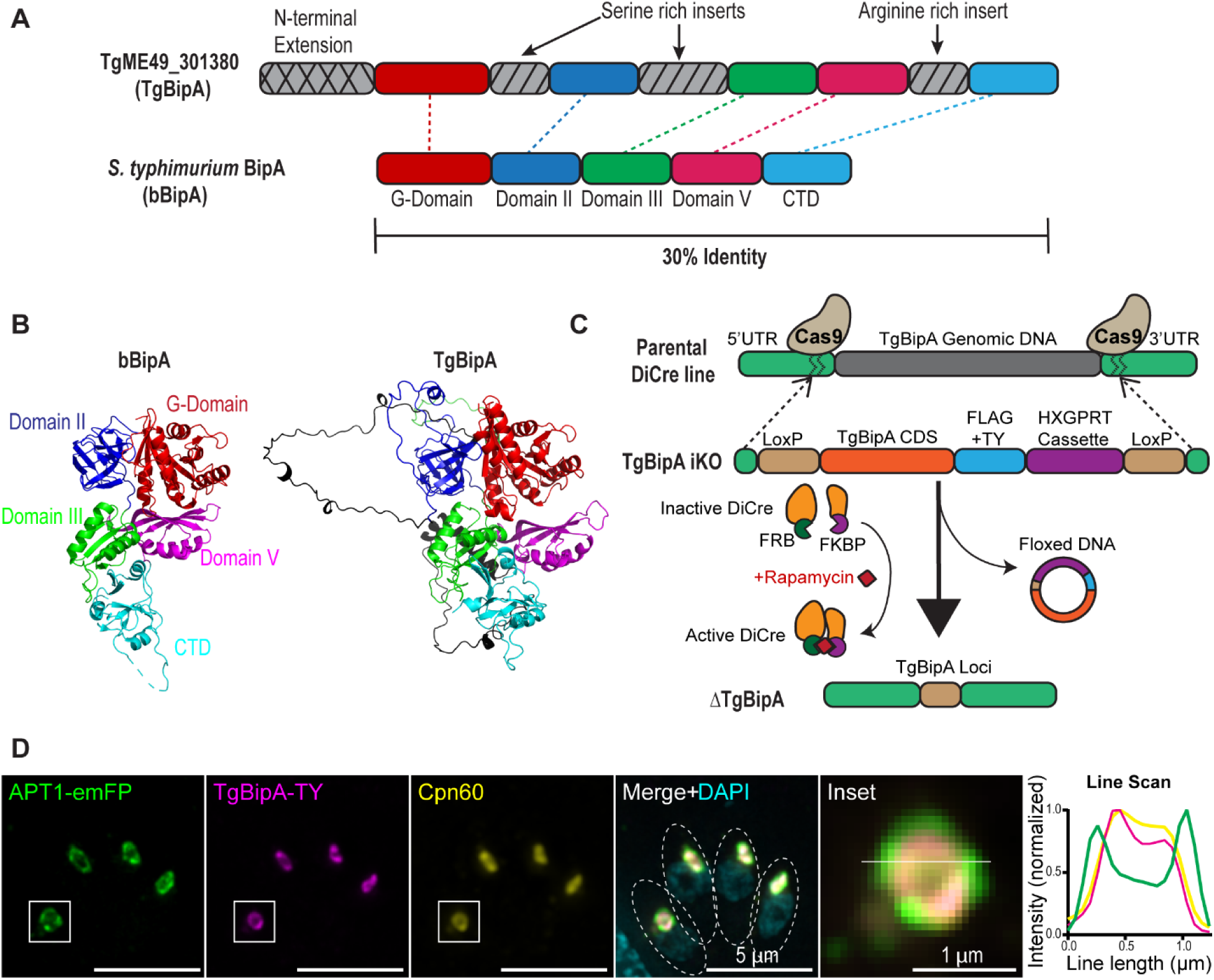
TgBipA is an apicoplast localized protein and homolog of BipA. A) Domain conservation between TgBipA and *S. typhimurium* BipA. TgBipA contains an N-terminal extension and three inserts not found in BipA. B) Comparison of *S. typhimurium* BipA protein structure (PDB 8EWH) and Alphafold 3 model prediction of TgBipA, minus the first 500 amino acids corresponding to the N-terminal extension. TgBipA unique inserts are in black. C) Schematic of TgBipA-iKO and TgBipA knockout (ΔTgBipA) strategy. In the RH_Ku80:Dicre:T2A:CAT parasite line, the endogenous TgBipA gene was replaced with LoxP flanked TgBipA coding sequence containing a FLAG and TY tag, followed by an HXGPRT drug selection cassette. Rapamycin treatment results in the activation of Cre recombinase and results in the excision of the floxed construct resulting in TgBipA deletion D) Immunofluorescence showing colocalization between TgBipA using anti-Ty antibody (magenta) and the apicoplast membrane marker APT1 fused to emFP (green), and the apicoplast lumen marker Cpn60 labeled using an anti-Cpn60 antibody (yellow). Merge+DAPI (cyan) shows a juxtanuclear position of the apicoplast. Dotted circles denote parasite outlines. Inset of apicoplast with line drawn indicating the region used to make line intensity profile which shows APT1 surrounding TgBipA and Cpn60 signals. Images are maximum intensity projections of deconvolved images. Scale bar = 5 µm. Inset scale bar = 1 µm.

To study the role of TgBipA, we created a TgBipA inducible knockout line, henceforth referred to as TgBipA-iKO, using the previously established DiCre system^56^. This line expresses Cre recombinase split into two components that are tagged with the rapamycin (rapa) binding peptides (FRB and FBKP). Rapa treatment induces dimerization to form an active Cre recombinase (Fig. 1C). Using CRISPR-Cas9, we replaced the endogenous TgBipA gene with the TgBipA coding sequence C-terminally tagged with Flag and Ty1 epitopes, and a drug selection cassette flanked by LoxP sites. Accurate integration into the TgBipA locus was confirmed by genomic PCR (Fig. S2A). This line was then modified to express the apicoplast membrane marker APT1 fused to EmeraldFP (EmFP) under the control of the APT1 promoter from the dispensable UPRT locus (referred to as TgBipA-iKO:APT1-EmFP hereafter) (Fig. S2B)^21,57^. To determine the localization of TgBipA, an immunofluorescence assay (IFA) was performed on this line using anti-Ty1 and anti-Cpn60 antibodies that recognize the TgBipA epitope tag and an apicoplast lumen protein, respectively^32^. We observed co-localization between TgBipA and Cpn60 signals, surrounded by the APT1-EmFP signal, showing that TgBipA localizes to the apicoplast lumen (Fig. 1D).

### TgBipA is essential to parasite survival

To determine the efficiency of TgBipA knockout after rapa treatment we activated the DiCre recombinase by treating parasites with 50nM rapa for 4 hours and then examined the TgBipA expression levels by western blot at 24-hour intervals starting at 48 hours post treatment. 48, 72, and 96 hours after rapa treatment there was 65%, 92%, and 97% decrease in TgBipA-TY protein levels respectively, relative to control (Fig. 2A, 2B and S3B). Consistent with this TgBipA protein levels were undetectable by immunofluorescence 72 hours after rapa treatment (Fig. 2C and Fig. S3A). To determine if TgBipA was essential for parasite survival, a plaque assay was performed using the DiCre parental line and TgBipA-iKO parasites treated with DMSO or rapa. TgBipA-iKO parasites treated with rapa failed to form plaques, indicating an essential role of TgBipA in the parasite lytic cycle (Fig. 2D).

**Fig. 2.**
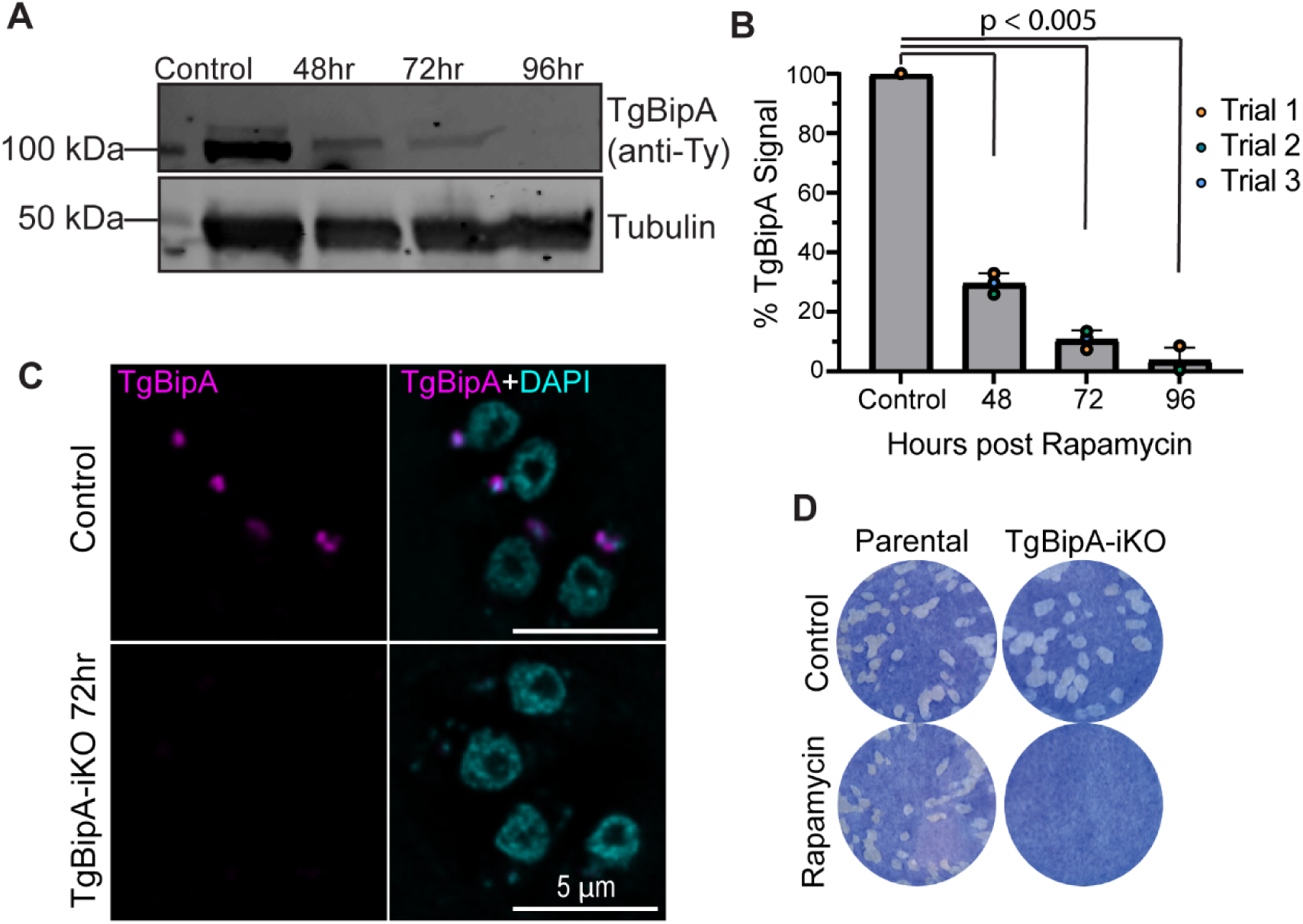
TgBipA is essential for parasite survival. A) Western blot with anti-TY (TgBipA) and anti-tubulin (loading control) antibodies on parasites lysates harvested at 48, 72, and 96 hours post rapamycin treatment, along with control. TgBipA runs at 110kDa, smaller than the 140 kDa predicted for the full-length protein suggesting TgBipA is proteolytically processed. B) TgBipA band intensities from (A) were quantified relative to tubulin loading control. Mean and individual values from 3 biological replicates were plotted. Significant p values are indicated (student’s paired t - test). C) IF of TgBipA-iKO parasites treated with DMSO (control) or rapamycin (KO) stained with anti-TY antibodies (TgBipA; magenta) and DAPI (nucleus) TgBipA is undetectable 72 hours after rapamycin treatment. Images are maximum intensity Z - projections of deconvolved images. Scale bar = 5 µm. D) Plaque assay of DiCre:T2A:CAT parental line and TgBipA-iKO line treated with DMSO or rapamycin prior to assay initiation. No plaques form upon TgBipA knockout indicating it is an essential protein.

### TgBipA is required for apicoplast maintenance

As TgBipA localizes to the apicoplast and is essential, we investigated if loss of TgBipA impacted apicoplast morphology by examining the localization of APT1 and Cpn60 in interphase parasites. In control parasites (TgBipA-iKO::APT1-EmFP; untreated), a clear ring-like structure of APT1 is observed around the Cpn60 and TgBipA signals (93 ± 2.9% of parasites) (Fig. 3A and 3B). 72 hours after rapa treatment, APT1 appeared ring-like in only 42 ± 7.5% of parasites. In 45 ± 2.8% of parasites, APT1 had a disordered signal, and no clear ring was visible. 13 ± 4.8% of parasites had multiple dim APT1 puncta. At the 96-hour time point, these percentages shifted to 33 ± 5.4% and 60 ± 4% of parasites for disordered and punctate, respectively. Additionally, apicoplast DNA could be visualized using DAPI in 85 ± 5.1% of controls compared to just 30 ± 7.7% rapa treated parasites grown for 96 hours after knockout (Fig. S3C). For localization of the lumen marker Cpn60, we see a similar trend to that observed for APT1. 97 ± 2.9% of untreated parasites showed a single bright punctate, consistent with the expected apicoplast lumen localization, this percentage dropped to 76 ± 9.5% 72 hours after knockout while 13 ± 7% contained faint Cpn60 puncta in addition to the apicoplast. 11 ± 7.3% did not contain a discernable apicoplast but multiple dimmer puncta were visible. 96 hours after rapa treatment, the Cpn60 localization is further disrupted with only 22% ± 11.7% of parasites containing a single Cpn60 puncta while 52 ± 11.7%, contained dim Cpn60 puncta (Fig. 3A and 3C). We did not observe any parasites completely lacking APT1 or Cpn60 signals. Notably, the disruption in Cpn60 appears to lag behind that of APT1, indicating temporal differences in how membrane and lumen proteins are disrupted after a TgBipA knockout.

**Fig. 3.**
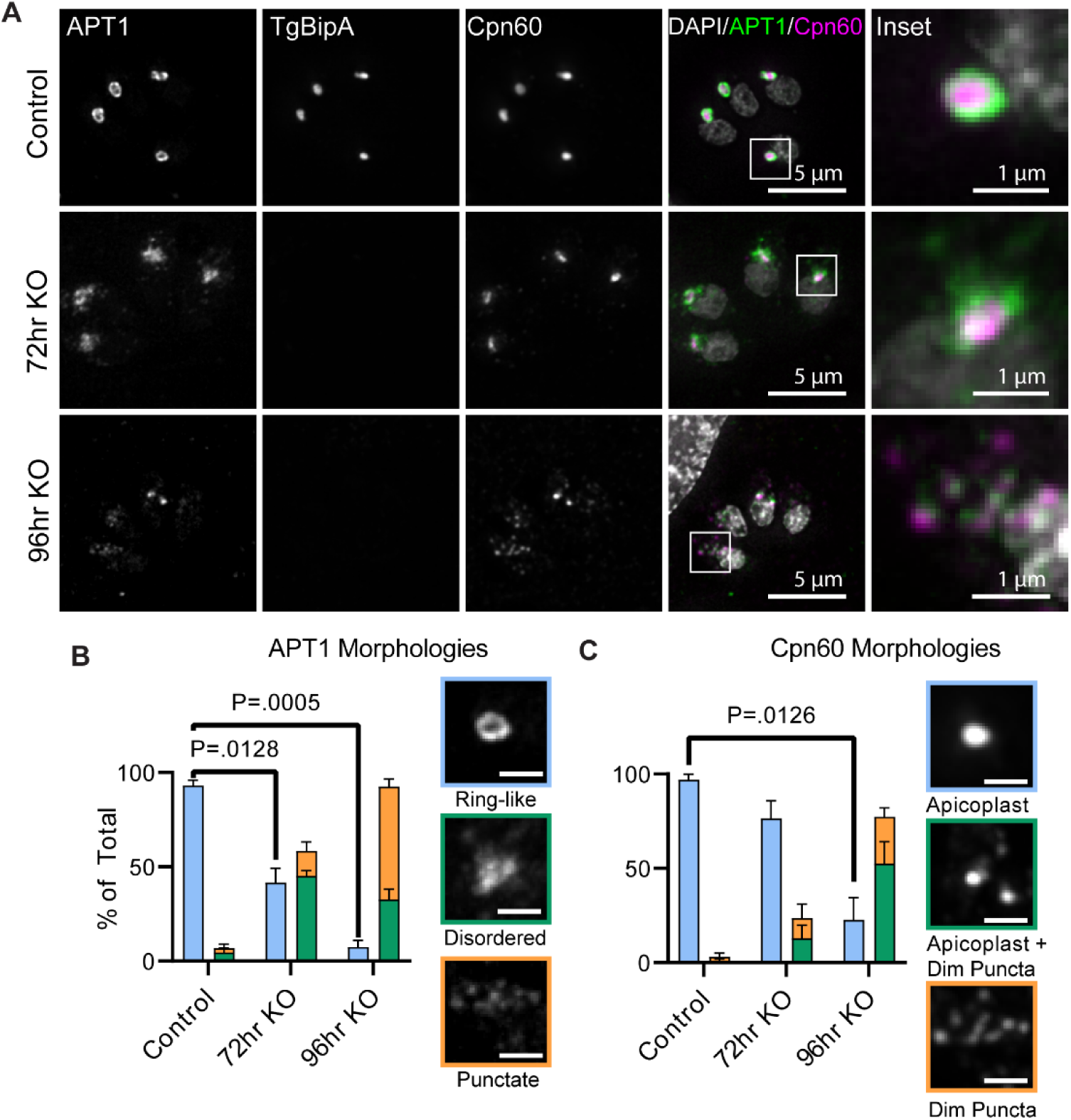
TgBipA KO disrupts apicoplast morphology. A) Immunofluorescence comparing TgBipA-iKO::APT1-GFP and ΔTgBipA:: APT1-GFP parasites 24 hours after invasion and 72 hours, and 96 hours after rapamycin treatment. TgBipA and Cpn60 visualized using anti-TY and anti-Cpn60 antibodies respectively. White boxes denote inset region. In 96hr KO images, the brightness was adjusted to show fainter punctate. Images are maximum intensity Z - projections of deconvolved images. Scale bar = 5 µm, inset = 1 µm. B) Quantification of APT1 and C) Cpn60 morphologies from time course as a percent of total. Data represents 3 independent experiments with n of at least 100 parasites. Examples of each phenotype shown in colored boxes matching phenotype bar chart color. Significant p - values from student’s paired t - test are indicated.

Loss of TgBipA disrupts apicoplast morphology such that the membrane labeled with APT1 could not be resolved as a distinct ring in diffraction limited images. To further investigate how TgBipA depletion alters apicoplast size and morphology, we performed 4x ultra-structure expansion microscopy^58,59^ and measured the area of the APT1 ring (Fig. 4A). The average expanded apicoplast area was reduced by ∼40% from 3 ± 0.75 µm^2^ in control parasites to 1.7 ± 0.39 µm^2^, 72 hours post treatment (Fig. 4B).

**Fig. 4.**
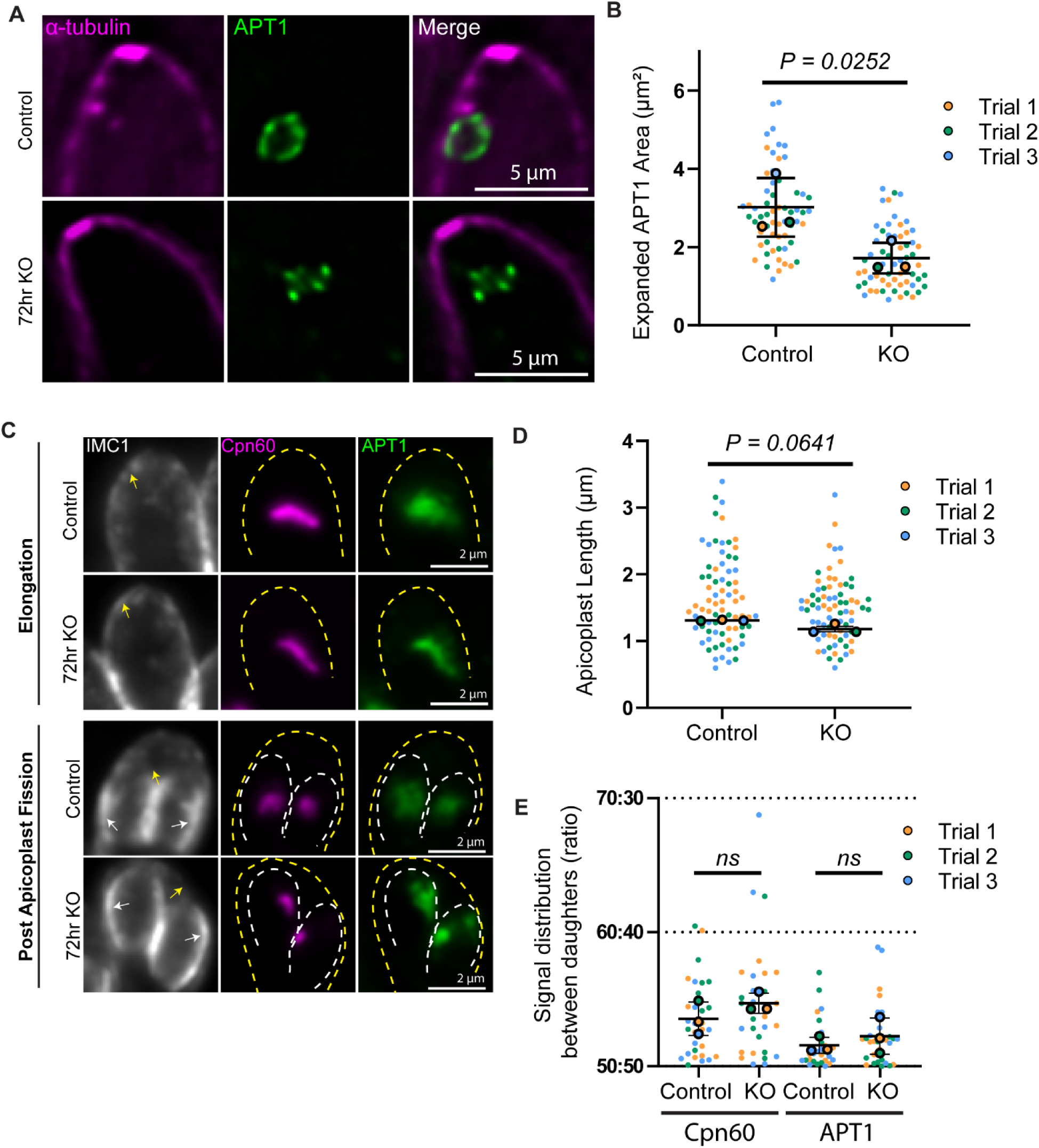
A) Expansion microscopy images of TgBipA-iKO::APT1-GFP and ΔTgBipA::APT1-GFP parasites. Samples were stained with anti-GFP and anti-tubulin antibodies. Images are single Z – slices of deconvolved images. Scale bar = 5 µm. B) Area measurements of expanded apicoplast using APT1 as a marker for apicoplast periphery, normalized to expansion factor. Mean area from each experiment is indicated with the large circles, individual values are indicated with small circles. Significant p - values from student’s paired t - test are indicated. Data from 3 independent experiments, n of 25. C) Apicoplast inheritance TgBipA-iKO::APT1-GFP and ΔTgBipA::APT1-GFP (72-hour time point) parasites. Apicoplast visualized through APT1-EmFP (green) and staining with anti-Cpn60 antibody (magenta). Periphery of mother (yellow arrows and dashed lines) and daughter (white arrows and dashed lines) parasites visualized using an antibody against inner membrane complex 1 (IMC1) (grey). Images are maximum intensity Z - projections of deconvolved images Scale bar = 2 µm. D) Graph showing lengths of apicoplast during elongation phase of division in TgBipA-iKO::APT1-GFP and ΔTgBipA::APT1-GFP parasites. Data from 3 independent trials, n of 20. p – values from student’s paired t – test are shown. Graph showing distribution of APT1 and Cpn60 signal in daughters in TgBipA-iKO::APT1-GFP and ΔTgBipA::APT1-GFP (72-hour time point) parasites. Data from 3 independent trials, n of 10 daughter pairs.

### Loss of TgBipA does not affect apicoplast inheritance

As we observed a defect in APT1 localization beginning 72 hours after TgBipA knockout, we next investigated whether this defect was due to a disruption of apicoplast inheritance. Apicoplast division is a multi-step process: early in cell division, the apicoplast elongates and then associates with the duplicated centrosomes in an ATG8 and actin dependent manner^31,60–65^. As the daughter cytoskeleton is constructed, the centrosomes and apicoplast ends migrate into the daughter parasites which results in the apicoplast forming a u-shaped structure^31^. DrpA-mediated apicoplast fission results in each daughter inheriting a single apicoplast^66^.

There were no differences in the length of elongated apicoplast in control and TgBipA knockout parasites, 1.64 ± 0.01 µm compared with 1.47 ± 0.09 µm for respectively) (Fig. 4C and 4D). Later in division cycle, the apicoplast was successfully inherited by daughter parasites. Fluorescent intensity measurements demonstrated that the Cpn60 and APT1 signals are approximately equal in daughter parasites in both control and knockout parasites (Fig. 4C and 4E).

### Loss of TgBipA results in the accumulation of APT1 vesicles

The apicoplast membrane appears less defined upon loss of TgBipA. To better understand this altered membrane morphology we performed live-cell imaging using the TgBipA-iKO:APT1-emFP line transiently expressing the apicoplast lumen marker FNR fused to RFP. In control interphase parasites, APT1 has a characteristic ring-like structure which surrounds FNR-RFP, with few observable APT1 vesicles (Fig. S4A and Video S1). At 72 hours post rapa treatment, we observed interphase parasites with a disordered APT1 signal surrounding the FNR-RFP signal, with an accumulation of APT1 vesicles surrounding these structures (Fig. S4B and Video S2). Thus, the “disordered” appearance of the apicoplast is due to a combination of a decrease in apicoplast size and accumulation of adjacent vesicles. Consistent with our fixed imaging, we did not observe localization defects in FNR 72 hours after knockout. However, beginning at 96 hours we do observe FNR vesicular structures in the apical region of the parasite which do not to colocalize with the APT1 vesicles (Fig. S4C and Video S3), consistent with fixed cell data (Fig. 3).

### Accumulated NEAT membrane vesicles have an ER origin

We next wanted to identify the origin of the apicoplast vesicles accumulating in the TgBipA KO. We postulated that these vesicles could be due to either an accumulation of ER-derived vesicles^19,67,68^, or a destabilization of the apicoplast leading to shedding of APT1-labelled membrane through an unknown pathway. To determine which was occurring, we expressed Halo tagged APT1 from the UPRT locus in the TgBipA-iKO line^31^ (TgBipA-iKO line::APT1-Halo) (Fig. S5A) and used a pulse-chase method to differentially label existing and newly synthesized APT1 protein with cell permeable HaloTag fluorescent ligands^31^. Differentiating between existing and newly synthesized proteins relies on saturation of existing halo epitopes with halo ligand. To confirm that this achieved in our experimental parameters, TgBipA-iKO line::APT1-Halo parasites were grown for 24 hours and then pulse labeled with TMR ligand for 30 minutes. Cells were washed and then immediately labeled with JF646 halo dye for 30 minutes, washed and immediately imaged (Fig. S5B). In all parasites, TMR strongly labelled the apicoplast, while JF646 was absent from the apicoplast, indicating that the TMR label saturates the Halo protein. Thus, in subsequent experiments, JF646 labeling is indicative of new protein synthesis.

This was a big paragraph … can we start a new one hereWith this system, we sought to determine the source of vesicles that accumulate in the cytosol after TgBipA disruption. 48 hours after rapa treatment, parasites were added to Mattek dishes, grown for 2 hours and then pulse labeled with TMR-Halo ligand, grown for a further 22 hours before pulse labeling with JF646, washed and imaged after an 8-hour outgrowth (Fig. 5A). APT1-TMR was predominantly localized to the apicoplast in both control and knockout conditions indicating APT1 protein is inherited by daughter apicoplast during division (Fig. 5B; green). There was significant variability in the localization of APT1-JF646. In some parasites, JF646 was almost exclusively localized to the apicoplast and there was significant overlap between the JF646 and TMR signals (Fig 5B; top panel shows an 80% overlap). In other parasites, the JF646 signal was visible in the ER or in cytosolic vesicles, indicative of protein that is newly synthesized and trafficking to the apicoplast (Fig. 5B; third panel 40% overlap). The mean overlap between TMR and JF646 in control parasites is 60 ± 8.67% (Fig. 5C). 80 hours after TgBipA knockout, the mean overlap between the two fluorophores was significantly reduced to 36 ± 7.1% and APT1-JF656 was visible in cytosolic vesicles and in the ER (Fig. 5B and 5C). The accumulation of APT1-JF646 vesicles in the TgBipA knockout, and the absence of APT1-TMR in vesicles, indicates that loss of TgBipA disrupts APT1 trafficking and leads to the accumulation of newly synthesized protein in vesicles in the cytosol. To validate this trafficking defect, we investigated if loss of TgBipA resulted in an accumulation of immature NEAT proteins^34,36^. Western blots were performed on controls samples and those grown for 48, 72 and 96 hours after rapa treatment using anti-Cpn60, anti-TY (TgBipA), and anti-tubulin as a loading control. In control and 48-hour samples, almost all Cpn60 is in the mature cleaved form (mCpn60) with virtually no uncleaved pre-mature Cpn60 (pCpn60) detectable. An accumulation of preCpn60 is observed for the 72- and 96-hour time points (Fig. S6A).

**Fig. 5.**
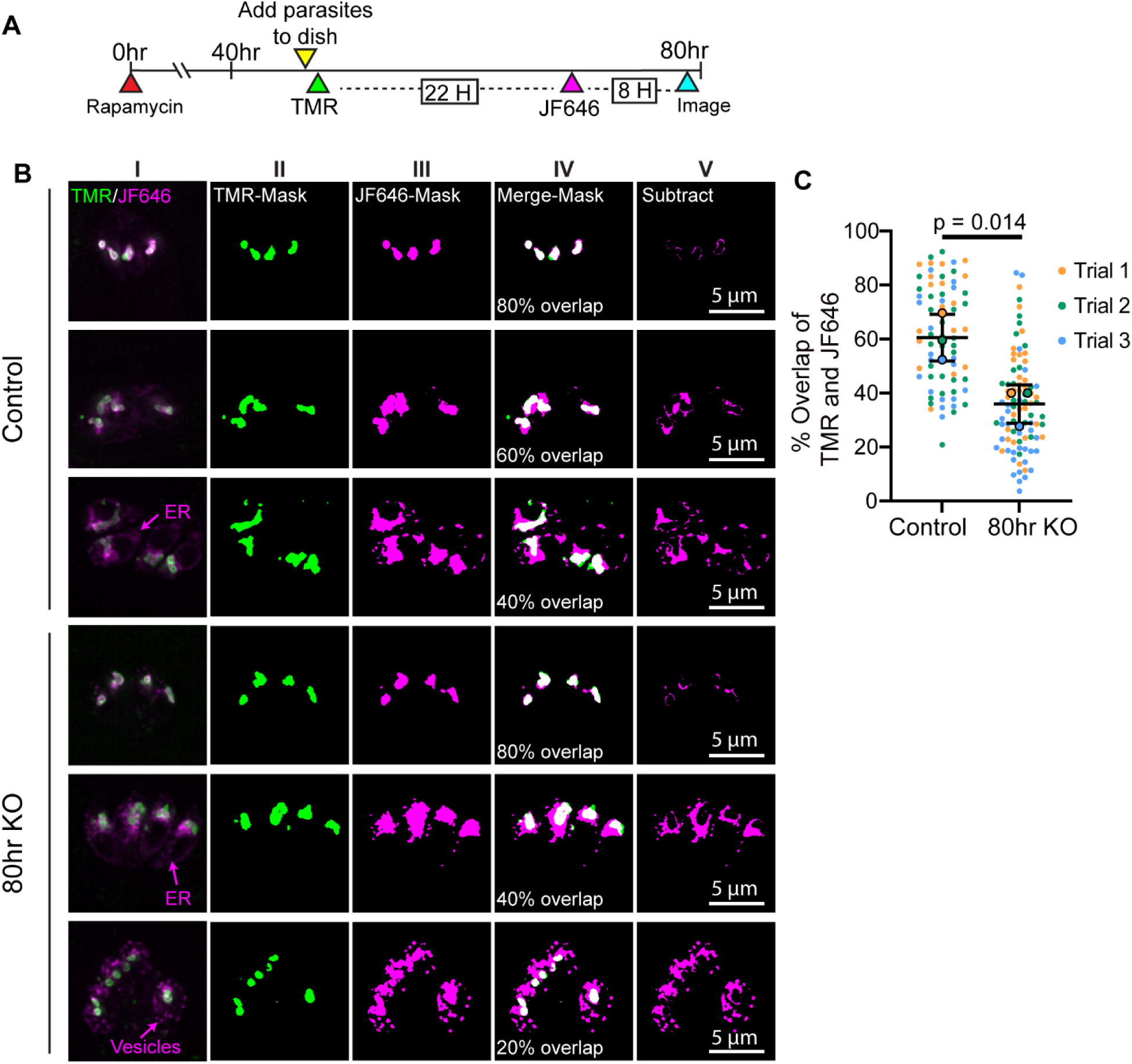
TgBipA KO leads to accumulation of ER derived vesicles labeled with NEAT membrane proteins. A) Timeline for pulse-chase assay to visualize APT1-Halo trafficking. B) TgBipA-iKO::APT1-Halo and ΔTgBipA::APT1-Halo parasites labeled with TMR and JF646 dyes as outlined in (A). In all parasites, Halo-TMR (green; pulse 1) had an apicoplast localization. JF646 (magenta; pulse 2) had a variable distribution and variability in overlap of JF646 and TMR dyes. Column I: are maximum intensity projections of deconvolved images. Columns II and III: TMR and JF646 images visualized as binary images. Column IV: Merge of binary TMR and JF646 images. Percent overlap is indicated. Column V: Signal from binary JF646 minus binary TMR signal (Subtract). Scale bar = 5 µm C) Percent of JF646 overlapping with TMR in images from B (see materials and methods for details). Mean overlap from each experiment is indicated with the large circles, individual values are indicated with small circles. Significant p - values from student’s paired t – test. Data from 3 independent trials, n of 20 - 30 vacuoles.

### Apicoplast translation inhibition mirrors TgBipA knockout phenotypes

All bacterial trGTPases interact with the ribosome including bBipA and thus it is possible that TgBipA might stably be associated with ribosomal components ^46,47,69^. Thus, we performed Flag immunoprecipitation on lysates from TgBipA-iKO parasites and RH parasites as a control to identify TgBipA interacting proteins. While we successfully purified TgBipA from iKO lysates (Fig. S7A), we did not identify any ribosomal components that were significantly enriched in this pulldown compared to controls (Table. S4). In the LC MS-MS analysis of TgBipA, all the identified peptide fragments corresponded to amino acid 413 or greater, and we speculate the N-terminus region likely contains the apicoplast targeting motif which is proteolytically processed during trafficking (Fig. S7B).

As an alternative approach to address a possible role for TgBipA in apicoplast translation, we compared the TgBipA KO phenotypes to parasites treated with 1 µM doxycycline (DOX). At this dosage, DOX specifically blocks apicoplast translation without inhibiting mitochondrial translation^39,40^. After 72 hours of continuous DOX treatment, we observed a disordered APT1 signal, along with an accumulation of APT1 and Cpn60 vesicles (Fig. 6A). At 96 hours of continuous treatment, APT1 and Cpn60 are found in punctate structures in the cytosol. Thus, DOX treatment and loss of TgBipA result in similar apicoplast morphology defects (Fig. 3). In addition, DOX treatment results in the accumulation of immature NEAT proteins, similar to the defects observed after TgBipA knockout (Fig. S6A)

**Fig. 6.**
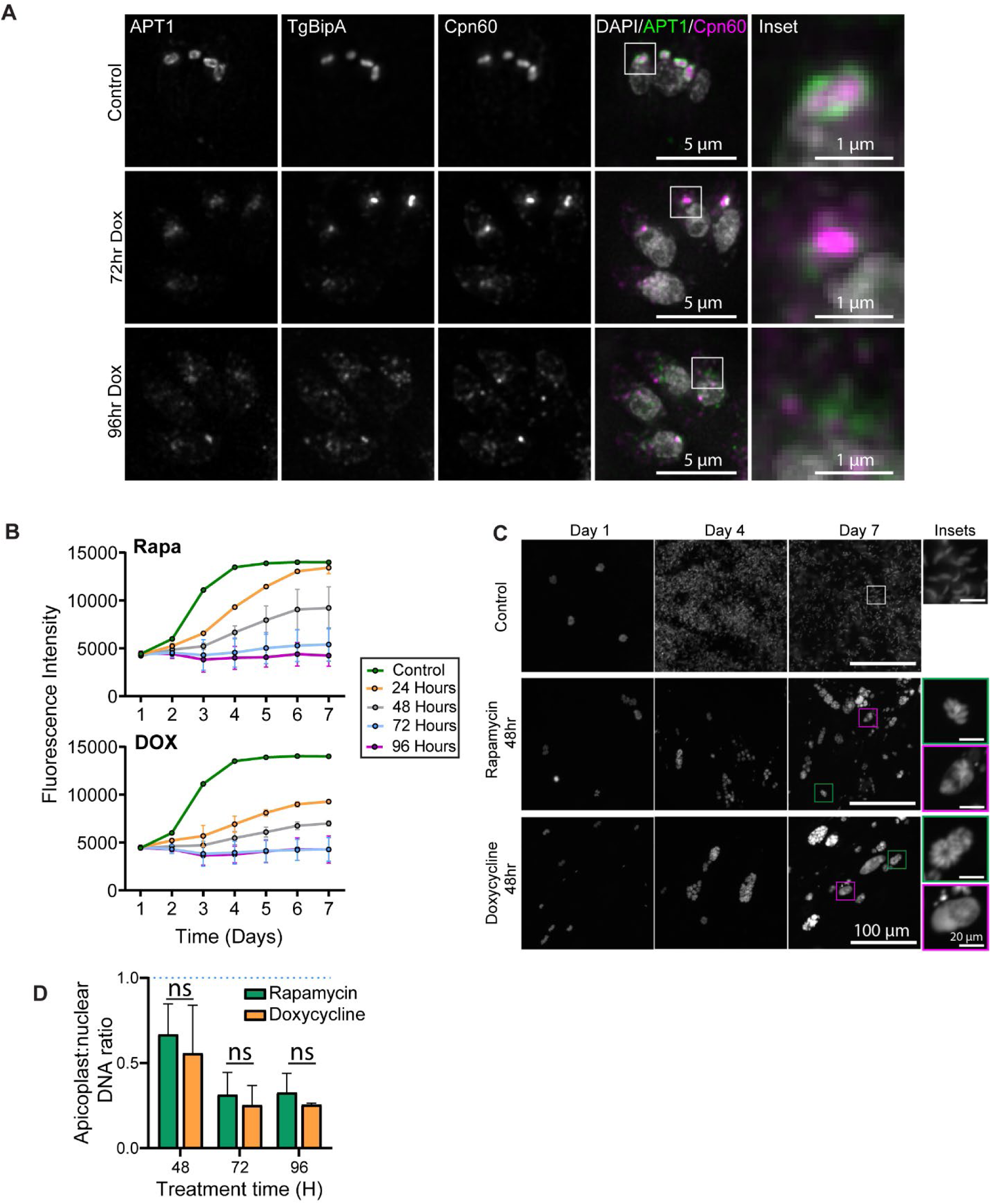
Doxycycline treated parasites and ΔTgBipA parasites display similar phenotypes. A) Immunofluorescence time course of TgBipA-iKO::APT1-EmFP parasites 24 hours post invasion. Untreated parasites were compared to parasites treated with DOX for 72 and 96 hours. TgBipA and Cpn60 visualized using anti-TY and anti-Cpn60 antibodies respectively. White boxes denote inset region. Brightness of 96 hr. KO images were adjusted to show fainter punctate signal. Scale bar = 5 µm, inset = 1 µm. B) Spectrophotometer fluorescence intensity readings of TgBipA-iKO:: GFP parasites were taken daily for 7 days. Growth rate of control parasites was compared to parasites treated with rapamycin, 24, to 96 hours prior to the start of the experiment or parasites grown continuously in doxycycline for 24-96 hours prior to the start of the experiment. Data from 2 independent experiments with technical triplicate. C) Images of wells from control and 48-hour rapamycin or DOX samples at day 1, 4 and 7. Green box denotes parasite vacuole with healthy cytosolic emFP signal, magenta boxes denote vacuole with emFP signal in the PV. Scale bar = 100 µm. Inset = 20 µm D) qPCR to determine relative copy numbers of apicoplast and nuclear genomes at control, 48, 72, or 96 hour time points. Blue dotted line represents control conditions. Performed in duplicate with technical triplicate. Student’s paired t – test comparing treatment conditions are shown.

### TgBipA KO and doxycycline treated parasites have similar growth rates

Next, we compared the impact of TgBipA knockout and DOX treatment on parasite growth. In the TgBipA-iKO parasite line, an EmeraldFP (EmFP) expression cassette was inserted into the UPRT locus (TgBipA-iKO::EmFP) (Fig. S5A). 24, 48, 72, and 96 hours prior to seeding, parasites were pulse treated with rapa or cultured continuously in DOX and then seeded into HFF monolayers in a 96 well plate. GFP fluorescence was measured with a spectrophotometer daily for 7 days and growth rates compared (Fig. 6B). In addition, the wells were imaged with epifluorescence microscopy to investigate parasite vacuole health (Fig. 6C and Fig. S8). In untreated parasites, we see an increase in fluorescence on days 1-4, followed by a plateau. This coincides with full lysis of the host cell monolayer observed via microscopy (Fig. 6C). For parasites treated 48 hours prior with rapa or continuously with DOX we see a ∼35% and ∼50% decrease in growth at day 7. When these wells were visualized by microscopy, we saw a mixture of healthy-looking vacuoles containing GFP positive parasites in a rosette pattern (Fig 6C; green inset box) and unhealthy vacuoles where GFP had leaked into the PV (Fig 6C; magenta inset box). Almost no growth is observed when the assay was started 72 and 96 hours after treatments (Fig. 6B). No treated parasites formed plaques in a plaque assay, which indicates that plaque assays are an imprecise method of assessing parasite growth (Fig. 6B and Fig. S7B). Thus, TgBipA KO and DOX treated parasites display similar growth kinetics, however TgBipA knockout growth defects lagged dox treatment, which we hypothesize is due to lag between the genomic excision of TgBipA and protein turnover.

### Loss of TgBipA results in apicoplast genome replication defects

One of the earliest markers of apicoplast translation inhibition after DOX or clindamycin treatment is a reduction in apicoplast genome copy number due to failure in apicoplast genome replication in the first lytic cycle^45,49^. To determine if this phenotype was observed in BipA knockouts, we used qPCR to determine the relative copy number of the apicoplast localized gene *TufA* to the nuclear gene *Act1* after rapa and DOX treatments^45,49,70^ (Fig. 6D). For each of the treated conditions, we saw a ∼50% reduction in apicoplast genome copy number at 48 hours post treatment, this was further reduced to ∼70% at 72 and 96 hours. Thus, loss of genome copy number is the first observable defect after loss of TgBipA or DOX treatment, occurring prior to defects in NEAT trafficking.

### TgBipA exhibits GTPase activity *in vitro*

As stated previously, the G-domain of TgBipA is 55% identical to that of its bacterial counterpart. Utilizing a malachite-green endpoint assay, we confirmed TgBipA’s ability to hydrolyze GTP suggesting it is a functional GTPase (Fig 7C). One of the recognizable features of any G-protein is the G1-motif, ^530^AHVDHGKT^537^ in TgBipA. In Ras, substitution of the lysine in this motif causes severely reduced GTPase activity because the protein cannot properly coordinate the guanine nucleotide or a Mg^2+^ in its active site generating an inactive GTPase^71^. A similar substitution in TgBipA, K536A, resulted in GTP hydrolysis rates that fell below the detection level of the malachite green assay (Fig. 7C).

**Fig. 7.**
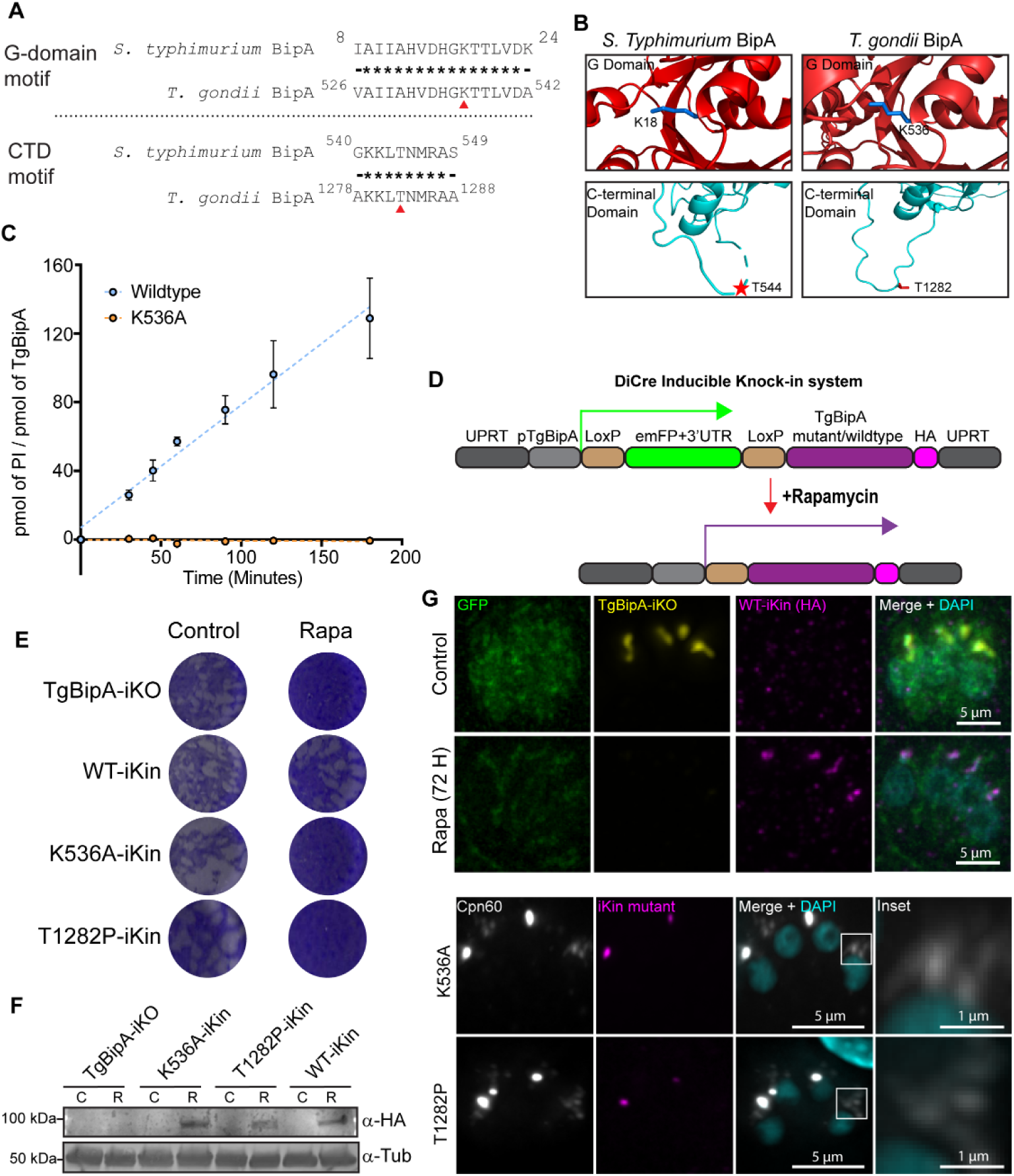
Conserved residues required for GTP hydrolysis and ribosome binding are essential for TgBipA function. A) Amino acid sequence comparison between the G1 motif and a C-terminal loop of bBipA and TgBipA. Conserved residues denoted by asterisk. Mutated residue denoted by red triangle. B) PDB and AlphaFold structures of G and CTD domains of bBipA and TgBipA respectively., K18/K536 in blue, and T543/T1282 in red. C) Absence of GTPase activity in the TgBipA K536A compared to wild type protein determined by a malachite green end point assay which detects levels of free phosphate in solution. Error bars represent standard deviation. D) Schematic of inducible knock-in (iKI) expression system. The UPRT gene was replaced with emFP which is expressed using the TgBipA promoter, flanked by LoxP sites. Upon rapamycin treatment emFP is excised and TgBipA wildtype (WT) or mutant are expressed. E) Plaque assays performed with TgBipA-iKO (parental), TgBipA WT-iKin, K536A-iKin, and T1282P-iKin parasite lines. F) Anti-HA western blot analysis of TgBipA-HA-iKin^WT/mut^ expression in control (C) parasites and 48 hours after rapamycin treatment (R). Tubulin was used as loading control. G) *Upper:* Immunofluorescence of WT-iKI rescue line in control and rapamycin treated conditions. TgBipA-iKO visualized by anti-TY antibody (yellow) and WT-iKI visualized by anti-HA antibody (magenta). *Lower:* K536A-iKI and T1282P-iKI lines 96 hours after rapamycin treatment. White box denotes inset region. Scale bar = 5 µm, Inset = 1 µm.

### Creation of a TgBipA knock-in/knock-out system

To investigate if GTPase activity was required for TgBipA function *in vivo*, we created a rapa inducible knock-out/knock-in (iKin) system to conditionally express TgBipA point mutants upon knockout of the wildtype protein (Fig. 7D and S9A). The UPRT gene was replaced with a LoxP inducible expression cassette. In untreated parasites the emFP coding sequence is transcribed under the TgBipA promoter and blocks the transcription of the downstream TgBipA gene. Upon rapa treatment, emFP is excised via Cre-recombinase and the TgBipA coding sequence (CDS) is repositioned downstream of the TgBipA promoter, inducing its expression^72^. This coincides with the simultaneous excision of wild type TgBipA gene from the TgBipA genomic locus. This approach allowed us to circumvent any potential dominant negative phenotypes that could arise from constitutive expression of the TgBipA mutant. We first investigated the efficiency of this Knock-out/Knock-in strategy by placing the HA-tagged wildtype TgBipA into knock-in locus (TgBipA^WT^-iKin). In untreated parasites, emFP is present in the cytosol and TgBipA-iKO (Ty-tagged) is expressed, while the TgBipA^WT^-iKin is not expressed (Fig. 7G). 72 hours after rapa treatment, GFP and TgBipA-iKO-Ty expression are not detectable, and TgBipA^WT^-iKin signal is observed at the apicoplast in 98% of parasites (Fig. 7G and S9B). Expression of TgBipA^WT^-iKin-HA from the UPRT locus rescued the growth defect observed in the TgBipA knockouts (Fig. 7E).

### TgBipA GTPase activity is required for parasite survival

To determine if GTP hydrolysis is necessary for TgBipA function, we tested if expression of TgBipA^K536A^-iKin could rescue the growth inhibition observed in the TgBipA knockout. We additionally tested a Thr1282Pro (T1282P) mutant, as the equivalent mutation in bBipA (T544P) has been shown to disrupt ribosome binding^69^. Anti-HA IF confirmed that both mutant proteins were localized to the apicoplast, indicating that these mutations did not disrupt trafficking to the apicoplast (Fig. 7F, Fig. S9A and S9C). However, unlike complementation with the WT, neither mutation could rescue the TgBipA KO growth defect (Fig. 7E). We next performed an anti-Cpn60 IFA to examine apicoplast morphology (Fig. 7G). In both mutants, we observe parasites lacking a Cpn60 puncta that instead contain vesicular Cpn60 signal (Fig. 7G Inset). Thus, expression of these mutations cannot overcome the apicoplast morphology defects of the knockout. Taken together, this data indicates that K536 and T1282 are essential residues for TgBipA function within the apicoplast.

## Discussion

The essentiality of the *T. gondii* apicoplast stems from its production of essential metabolites. Since most proteins required for these processes are nuclear encoded and trafficked to the apicoplast, the metabolic pathways are dependent on accurate trafficking of NEAT proteins from the ER, protein translocation across four apicoplast membranes, and chaperones and proteases required for protein maturation in the apicoplast lumen. In addition, two proteins encoded by the apicoplast genome, SufB and ClpC, are thought to contribute to metabolism, although their mechanism of action is unknown. Synthesis of these proteins relies on apicoplast ribosomal components, the majority of which are also encoded by the apicoplast. Here we describe an apicoplast localized GTPase, TgBipA that is essential for parasite fitness and maintenance of the apicoplast. Loss of this protein affects NEAT protein trafficking and maintenance of the apicoplast genome and therefore is essential for apicoplast function.

### The GTPase Activity of TgBipA is Essential

TgBipA is a homolog of the bacterial translational GTPase BipA^73^. Amino acid sequence comparison as well as an AlphaFold model of the protein indicates that the core regions of these proteins are analogous consisting of a GTPase domain, an OB domain, two alpha beta domains and the C-terminal domain, its novel fold characteristic of BipA family members^74^. Unique to TgBipA are three very large IDRs, distinctive in that two are serine rich and the third arginine rich. Their large size and propensity to undergo post-translational modifications suggest that TgBipA actively participates in diverse functions within the apicoplast^75^. Similar to its bacterial counterpart, TgBipA is an active GTPase. Mutating the conserved lysine in the GTP binding pocket of TgBipA to alanine (TgBipA K536A) results in a loss of GTP hydrolysis *in vitro*. TgBipA K536A cannot rescue a wild type TgBipA knockout, indicating that GTPase activity is critical for the function of the protein. In addition, like its bacterial counterpart, TgBipA has low intrinsic GTPase activity. Whether like prokaryotic trGTPases, the best known example being EF-G, TgBipA’s hydrolysis rate is significantly enhanced by partner binding, remains to be determined^76^.

TgBipA has an unusually long N-terminal extension of 500 amino acids. However, no canonical signal peptide or apicoplast targeting motif could be identified in this sequence. While there is no predicted targeting motif, transit peptides in *T. gondii* are typically enriched in serine residues^77^, and the TgBipA N-terminal extension is 28% serine which likely facilitates apicoplast trafficking. When mass spectrometry analysis of affinity purified TgBipA was performed, peptides corresponding to the first 412 amino acids were consistently missing from our analysis, indicating this sequence corresponds to the apicoplast targeting motif that is removed after import to the apicoplast.

### TgBipA is essential for apicoplast maintenance and NEAT protein trafficking

The first defect we observed following loss of TgBipA is a decrease in apicoplast genome copy number, similar to what is observed when apicoplast translation is inhibited using antibiotics (Figure 8, 48 hours)^45,49^. The link between apicoplast translation and genome replication is not understood, as none of the apicoplast encoded genes appear to have a direct role in replication. The two apicoplast encoded proteins not involved in translation, ClpC and SufB, have not been characterized but it has been speculated that these proteins have an indirect role in genome replication^23,36,37^. ClpC is predicted to be a homolog of a chloroplast ATP dependent unfoldase^36^, which in chloroplasts, interacts with the translocon at the inner chloroplast membrane (Tic) complex, and is thought to facilitate protein import^36,37,78,79^. Within apicomplexans, homologs of the chloroplast Tic-Toc complex localize to the innermost apicoplast membranes and are required for NEAT import^28,29,80^. Thus, it is possible that the import of DNA replication machinery such as the DNA-binding protein HU, involved in initiation of replication in other systems^81^, and DNA gyrase, that plays a role in DNA unwinding during DNA replication^82^, could be disrupted by an inhibition of apicoplast translation. However, our data argues against this hypothesis as we observed genome loss prior to the observation of defects in NEAT protein trafficking. Thus, an alternative hypothesis is that the iron-sulfur clusters provided by the SufB containing pathway may be required as cofactors for DNA replication enzymes^23,83^, and the loss of these cofactors results in defects in genome replication.

**Fig. 8.**
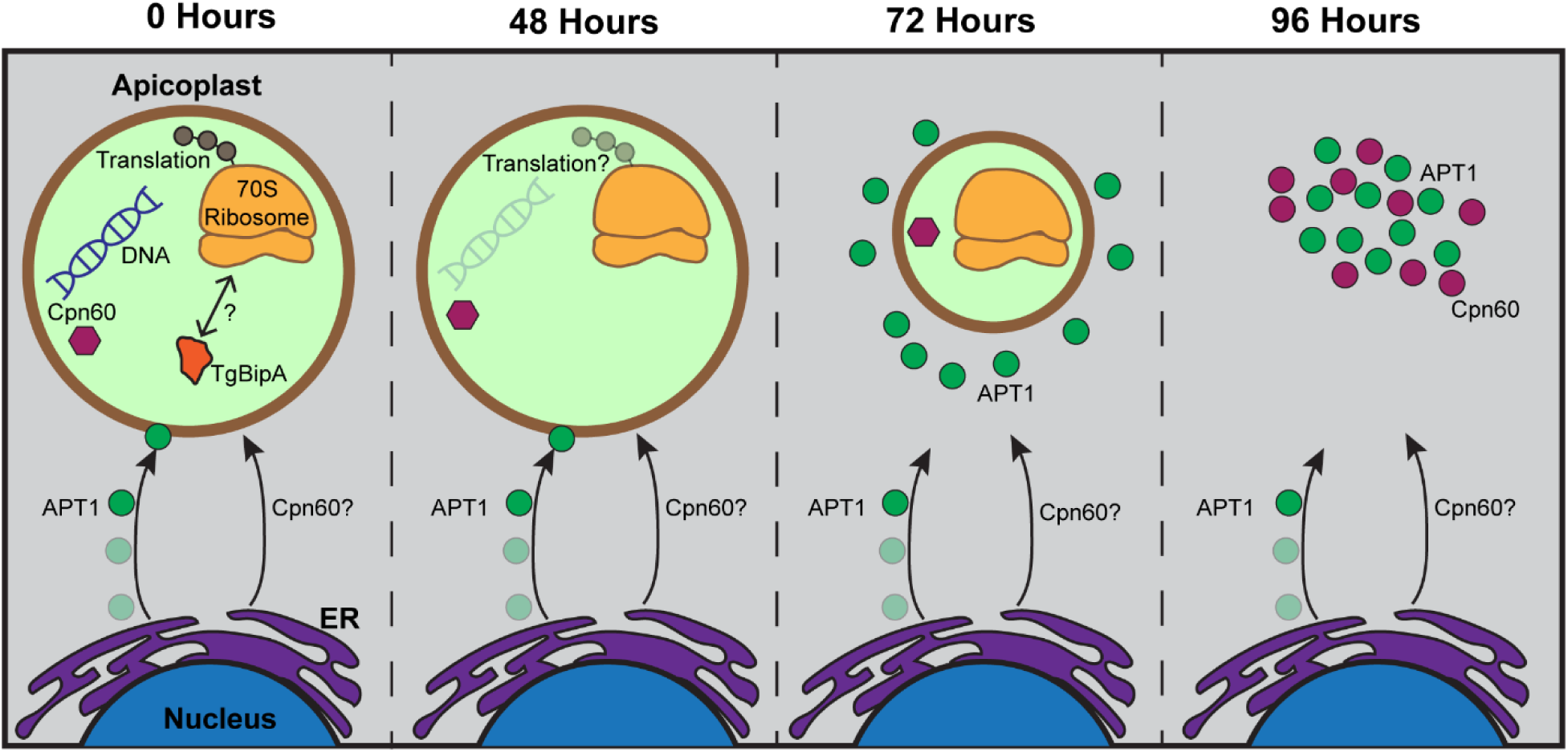
Working model for loss of apicoplast following TgBipA knockout. 48 hours after TgBipA genomic knockout, there is a reduction in apicoplast genome copy number. 72 hours after knockout there is a loss of APT1 trafficking and APT1-positive vesicles accumulate in the cytosol. There is a reduction in apicoplast size. After 96 hours, there is a loss of an observable apicoplast structure. APT1 and Cpn60 vesicles accumulate at the parasite’s apical end.

Following the loss of genome replication, we observed the accumulation of ER derived, APT1-labeled vesicles adjacent to the apicoplast and a decrease in apicoplast size (Fig. 8, 72 hours) as demonstrated through fixed cell (Fig. 3) and live cell (Video S1-3) imaging and pulse-chase assays (Fig. 5). It has previously been proposed that fusion of ER derived vesicles with the apicoplast would provide the additional lipids necessary for apicoplast expansion prior to division^25,31^. The simultaneous accumulation of vesicles and decrease in apicoplast size is consistent with this hypothesis. The pulse-chase assays show that APT1 protein is inherited by each apicoplast during division so the disruption in trafficking likely leads to a progressive decrease in protein levels through successive rounds of division. Phenotypic consequences of these trafficking defects would not be observed until membrane protein levels drops below a certain threshold and could provide an explanation for the delayed growth defects observed in *ΔTgBipA* parasites and after DOX treatment. As an example, APT1 is an apicoplast phosphate translocator and is responsible for the import of precursor molecules for fatty acid synthesis and thus essential for the apicoplast localized FASII pathway^10,57^. Thus, we speculate that a reduction in APT1 trafficking could indirectly contribute to the reduction in apicoplast size by reducing availability of fatty acids for membrane biogenesis.

24 hours after APT1-labeled vesicles are observed, Cpn60 containing vesicles began to form, and this is accompanied by loss of the apicoplast (Fig. 8, 96 hours). While it has previously been reported that membrane proteins traffic to the apicoplast in ER-derived vesicles, the pathway for NEAT lumen protein trafficking is not clear, as vesicles containing lumen proteins are not observed in healthy, wildtype parasites^19,30,31^. Alternative trafficking mechanisms have been proposed including ribosome docking on the apicoplast outer membrane^23^ or rapid transport in vesicles that is difficult to visualize experimentally^30^. Regardless of the mechanism in wildtype parasites, in *ΔTgBipA* parasites and parasites engineered to survive without the apicoplast^84^ Cpn60 and FNR are found in separate vesicles populations that accumulate in the apical region of the parasite. Given that disruption in lumen protein trafficking occurs ∼48 hours after the reduction in apicoplast genome copy number, this phenotype is likely a downstream, rather than direct, consequence of the TgBipA knockout.

Delayed death of *T. gondii* parasites following perturbation of the apicoplast has been well documented^39,42,45,49,50^. It is typically described as parasite death occurring in the second intracellular growth cycle (∼48-96 hours), with no growth defects observed in the first intracellular growth cycle (first ∼48 hours). An extensive time course carried out over the course of 7 days showed that ∼48 hours after loss of TgBipA and DOX treatment, parasite growth rates were reduced compared to control (Fig. 6). However, a small percentage of parasites remained viable for up to 10 days. It is unclear what accounts for this large variability in the kinetics of parasite death, but this observation demonstrates that death after apicoplast translation inhibition is more delayed than previously appreciated and underscores that translation inhibitors do not demonstrate the optimum killing dynamics for anti-parasitic’s.

In conclusion, this study characterizes a new apicoplast localized GTPase enzyme TgBipA. Loss of this protein results in a decrease in genome copy number, disruption of apicoplast morphology and trafficking of membrane-associated NEAT proteins, ultimately resulting in parasite death. These phenotypes are also observed when apicoplast translation is inhibited, however, despite our efforts we could not definitely determine if TgBipA has a direct role in translation, analogous to its bacterial counterpart. Future work will focus on identifying TgBipA’s mechanism of action at the apicoplast.

## Materials and Methods

### Cell Culture

Human foreskin fibroblasts (HFFs) were grown to confluency in Dulbecco’s modified Eagle’s medium (DMEM) (Fisher Scientific: Catalog#:11-965-092) supplemented with 1 x antibiotic/antimycotic (anti/anti), 1mM Sodium pyruvate and 10% heat-inactivated fetal bovine serum (FBS) at 37°C with 5% CO_2_. When cells were confluent, media was replaced with DMEM supplemented with 1x anti/anti and 1% heat-inactivated FBS. *T. gondii* parasite lines were continuously passaged in HFFs grown in low serum media.

### Drug treatments

For rapamycin treatment, 10mM Rapamycin (Invitrogen: Catalog#: tlrl-rap-5) in DMSO was diluted 1:100 in molecular biology grade water (Cytiva: Catalog#: SH30538.02) and added to DMEM at 1:2000 for a final concentration of 50nM. Parasites were grown in HFFs containing rapamycin media for 4 hours before washout. The start time for treatment is considered hour 0 for knockout.

For doxycycline treatment, 10mM doxycycline (Fisher BP26531) in DMSO was diluted 1:10 in water and used at a 1:1000 dilution for a final treatment concentration of 1µM. Parasites were continuously cultured in doxycycline containing media for time points indicated.

For parasite transfections which disrupt the UPRT loci, normal growth media was replaced with media containing 5’-fluo-2’-deoxyuridine (FUDR, Sigma Catalog#: F0503)) (10µM) 24 hours after transfection.

### Parasite Transections

All transient and stable parasite transfections were performed using previously described protocols^85–87^. Briefly, for transient transfections, 25 µg of plasmid DNA was transfected into 1 × 10^7^ parasites via electroporation. These parasites were grown overnight on a MatTek dishes or 6 well plates containing confluent HFF monolayers (Mattek Catalog#: P35G-1.5-14-C). To modify genomic loci using CrispR-Cas9, 5 µg of plasmid DNA containing Cas9 expression cassette and guide RNAs and homologous recombination oligomer (HR oligo) produced by PCR (50 ul) were transfected in and parasites grown overnight before switching to drug containing media. Parasites were subcloned by diluting to 100 parasites/15ml media and plating 1 parasite per well into a 96 well (Corning Catalog#: 3585) containing confluent HFF monolayer. Single plaques were scraped and plated into 8-well (ThermoFisher Catalog#: 155409) and 12-well (Corning Catalog#: 3512) plates. 8-well plates were imaged via microscopy to look for successful expression of tagged protein. Positive lines were confirmed via genomic PCR.

### Creation of plasmids and parasite lines

All primers, plasmids, and antibodies in this study can be found in Table S1, S2, and S3. For all Gibson assembly reactions we used a custom master mix using 160 µL of 5x isothermal buffer (500mM Tris-HCl, pH 7.5, 50mM MgCl_2_, 1mM dNTPs (each dNTP), 50 mM Dithiothreitol, 0.75g PEG 8000, 0.05 M β-NAD, raised to 3 mL in molecular biology grade water), 3.2 µL T5 exonuclease (Catalog#: NEB M0663S), 10 µL Phusion DNA polymerase (Catalog#: NEB M0530S), 80 µL Taq DNA ligase (Catalog#: NEB M0208L), in 346.8 µL Molecular biology grade water. This makes 40 −15 µL Gibson reactions.

#### Creation of TgBipA-iKO parasite line

Insertion of BipA gRNA into pU6 Universal plasmid. Complementary oligos containing 5’ and 3’ guide RNA sequences were reconstituted to a concentration of 200uM in molecular biology grade water, oligos were phosphorylated with PNK and then annealed by heating to 95C and then cooling at a rate of 0.1°C per second. Duplexed oligos were ligated to pU6 universal plasmid (pU6-Universal was a gift from Sebastian Lourido, AddGene plasmid # 52694), digested with BsaI and gel purified using T4 DNA ligase (New England biolabs) as per manufacturer’s instructions. Plasmid was transformed into chemically competent NEB5alpha. The sequence of positive clones were verified using whole plasmid sequencing (Plasmidsaurus). To create a plasmid containing both 5’ and 3’ BipA guide RNA’s, the pU6-5’ BipA plasmid was linearized with KpnI. The 3’gRNA, scaffold and promoter were amplified by PCR using KpnI-pU6promF and ptub-KpnI-R and then digested with KpnI enzyme. The linearized plasmid and digested PCR product were ligated using T4DNA ligase (NEB) as per manufacturer’s instructions. The sequence of positive clones was verified using whole plasmid sequencing (Plasmidsaurus).

To create the template for the HR oligo that contains the TgBipA coding sequencing containing Ty1 and Flag epitopes we first cloned the TgBipA coding sequencing into the pFastBac plasmid (unpublished plasmid). Due the size of the gene the coding sequence (CDS) was amplified with a series of nested primers (cBipA primers; Table S1). Fragments were then annealed to the linearized pFastBac plasmid containing a Flag epitope using Gibson assembly (unpublished plasmid). The BipA-Flag was then subcloned into a plasmid containing a LoxP site downstream of the HXGPRT selection cassette (Heaslip et al 2016), as illustrated in Fig. 1C. This plasmid (pTKOII-MyoF-LoxP) was digested with AvrII and AflII and plasmid backbone gel purified. TgBipA-Flag was amplified using primers BipA_CreLoxP and Flag-3xTy-R. Note the forward primer contains a LoxP site upstream of the BipA start ATG. 3xTy1 was amplified using primers 3xTy1-F and 3xTy1-R. Plasmid backbone and both PCRs were annealed using Gibson assembly as described above. Plasmid was transformed into chemically competent NEB5alpha. The sequence of positive clones was verified using whole plasmid sequencing (Plasmidsaurus).

#### Creation of pTub-emFP-UPRT parasite line

In the TgBipA-iKO:pTub-emFP-UPRT line, emFP is expressed from the UPRT loci under the control of the tubulin promoter. To create this line, the 5’UPRT-pAPT1-APT1-emGFP-3’UPRT plasmid (Table S2)^31^ was digested with AvrII and AflII to remove pAPT1-APT1-emGFP and the linearized backbone was gel purified following manufacturer’s instructions (Qiagen Catalog#: 28706). pTub-emFP-3’DHFR was amplified via PCR (unpublished plasmid, Table S3) with AvrII and AflII overhangs and gel purified and put into the UPRT backbone through Gibson assembly following manufacturer’s instructions (New England Biolabs Catalog#: M5520AA2). This plasmid was transformed into NEBDH5α cells. Positive clones verified through whole plasmid sequencing.

#### Stable transfection and integration of genomic construct into the UPRT loci

To create all HR oligo’s for transfection into the UPRT loci, a 50 µl PCR reaction was performed using UPRT F and UPRT R primers, Plasmid DNA template (Table S2), and a Q5 high fidelity DNA polymerase (New England Biolabs Catalog#: M0493L). This PCR product was transfected into 1×10^7^ TgBipA-iKO parasites along with 5µg of pSag1::Cas9::U6::sgUPRT plasmid (AddGene plasmid # 54467). 24 hours after transfection, media was replaced with media containing 10µM FUDR to positively select for UPRT loci disrupted parasites^88^. Positive clones were selected and confirmed for successful integration via genomic PCR using UPRT primers (Fig. S2B and Table S1 and S2)

#### Creation of TgBipA:APT1-emFP-UPRT and TgBipA:APT1-Halo-UPRT parasite lines

The plasmids used to make these lines was previously described in Devarakonda et al., 2023. Transfection into the UPRT loci was performed as described above.

#### Creation of TgBipA Inducible knock-in wildtype parasite line

For the TgBipA inducible knock-in (iKI) lines, mutant or wild type TgBipA is inducibly expressed from the UPRT loci using pTgBipA promoter and 3’UPRT UTR. To create this line, TgBipA CDS was PCR amplified from the pTKOII_TgBipA_CDSFlag_LoxP plasmid with overhangs to an HA gblock containing an NheI cut site and overhangs into the pTgBipA promoter with an NheI cut site. pTgBipA promoter was PCR amplified off of RH gDNA with TgBipA CDS and 5’UPRT overhangs containing NheI and NdeI cut sites. 3xHA-stop gblock with TgBipA CDS and 3’UPRT overhangs were ordered from IDT as forward and reverse ultramers and annealed together. All products were gel purified following the manufacturer’s instructions. 5’UPRT-pAPT1-APT1-emGFP-3’UTR was digested with AvrII and AflII to remove pAPT1-APT1-emGFP and the linearized backbone was gel purified following manufacturer’s instructions. Ligation of PCR products was performed via Giboson assembly to create 5’UPRT_pTgBipA_TgBipA CDS-3xHA_3’UPRT and transformed into NEBDH5α cells. Positive clones verified through whole plasmid sequencing. emFP-3’DHFR was amplified off of pTub-emFP-3’DHFR plasmid via PCR using ultramers with LoxP overhangs and homology to pTgBipA promoter and TgBipA CDS, and gel purified. 5’UPRT_pTgBipA_TgBipA CDS-3xHA_3’UPRT was digested with NdeI and gel purified. LoxP-emFP-LoxP was ligated into 5’UPRT_pTgBipA_TgBipA CDS-3xHA_3’UPRT backbone using Gibson assembly to create 5’UPRT_pTgBipA_Loxp-emFP-3’DHFR-Loxp-TgBipA CDS-3xHA_3’UPRT. This plasmid was transformed into NEBDH5α cells and positive clones verified through whole plasmid sequencing.

#### Creation of TgBipA^K536A^-iKin and TgBipA^T1282P^-iKin parasite lines

For creating inducible knock in rescue lines TgBipA^K536A^-iKin and TgBipA^T1282P^-iKin, 3’-DHFR-Loxp-TgBipA CDS with emFP and 3’UPRT overhangs was amplified off of 5’UPRT_pTgBipA_Loxp-emFP-3’DHFR-Loxp-TgBipA CDS-3xHA_3’UPRT in two pieces, with overlap at the mutation site using TgBipA K536A Mutant Fwrd/Rev (Table. S1) primers to create a base mutation resulting in a K/A amino acid change, and TgBipA T1282P mutant Fwrd/Rev (Table. S1) primers to make the T/P amino acid mutation, and PCR products were gel purified. 5’UPRT_pTgBipA_Loxp-emFP-3’DHFR-Loxp-TgBipA CDS-3xHA_3’UPRT was digested with AflII and HindIII and gel purified to remove 3’DHFR-Loxp-TgBipA CDS-3xHA. PCR products were inserted into the backbone using Gibson assembly and transformed into NEBDH5α cells and positive clones verified through whole plasmid sequencing. Constructs were transfected into UPRT loci as described above.

### qPCR

Parasite genomic DNA was extracted using Qiagen DNeasy Tissue kit (Catalog#: 69504) according to manufacturer’s instructions. qPCR was performed using iTaq universal SYBR green supermix (Biorad Catalog#: 1725121) on a Biorad CFX96 Real-Time PCR machine using primers in Table S1. Relative DNA quantity was determined using the Pfaffl method (Pfaffl 2001).

qPCR run information: Following an initial incubation at 95°C for 3 minutes, 40 cycles of 95°C for 10 seconds, 55°C for 20 seconds, 72°C for 20 seconds were performed.

### Fluorescence growth assay

Freshly lysed control and doxycycline or rapamycin treated parasites were diluted such that 6000 parasites were added to each well in Gibco fluorobrite DMEM (Catalog#: A1896701) supplemented with 1x anti/anti, 1% heat-inactivated FBS, and 4mM L-Glutamine. For doxycycline treated samples, assay was performed with media containing 1uM DOX. Parasites were added to Greiner 96 well plates (Catalog#: 655097) containing confluent HFF monolayer, and GFP fluorescence was measured daily using a Molecular Devices SpectraMax i3x. Each reported reading was an average of 21 measurements were taken per well.

### Western Blot

Parasites were syringe released, counted, and 6 × 10^7^ parasites were centrifuged at 1260g for 4 min. Cells were resuspended in 1xPBS to 2 × 10^7^ parasites/32ul, added to 8 µL 10x sample buffer (0.6M Tris-HCl pH 6.8, 700mM SDS, 20 % glycerol, 0.75mM bromophenol blue, 2.5mM 2-methcapethanol, 25mM Dithiothreitol, raised to 20 mL with water) for a final concentration of 2x sample buffer and boiled for 10 minutes at 95C. Samples equivalent to 2 × 10^7^ parasites were run on Bio-Rad 4-20% gradient gels (Catalog#: 4561094) and transferred to a nitrocellulose membrane. Membranes were blocked in blocking buffer (2% non-fat milk in 1xTBST (0.1M Tris-base pH 7.4, 10% Tween-20, 0.15M NaCl, in water) for 2 hours, and incubated with primary antibodies according to Table S3, diluted in blocking buffer. Membrane was washed three times for 10 minutes each in 1xTBST and incubated with secondary antibodies according to Table S3. Blot was washed and imaged on a LiCor Odyssey Fc with pierce ECL western blotting substrate (ThermoFisher Cat #: 32209), or on an Odyssey CLx.

### Plaque assays

To determine if TgBipA is essential for parasite viability, 200 TgBipA-iKO or RH:Dicre:T2A:CAT parasites treated 72 hrs. prior with rapamycin or an equivalent volume of DMSO were plated onto a six-well plate containing a confluent HFF monolayer and grown undisturbed for 7 days. Wells were fixed with methanol at −20°C for 15 minutes and then stained with Coomassie (53% H_2_O, 40% methanol, 7% acetic acid, 300µM Brilliant blue R (Sigma-Aldrich B7920)) for 2 hours and then destained (53% H_2_O, 40% methanol, 7% acetic acid). TgBipA mutant rescue plaque assays and doxycycline time course plaque assays were performed as above in a 12 well.

### Pulse-chase assay

For Halo-ligand labeling of APT1-Halo, parasites in a monolayer of HFFs in a MatTek dish were incubated with halo dyes as outlined in main text using TMR (Promega G8252) and Janelia fluor (Jf646) (Promega GA1120) at a final concentration of 1µM in DMEM for 30 minutes. Cells were washed 3x times in DMEM and imaged in fluorobrite.

### Fluorescence Microscopy

Imaging was performed on two microscopes: a DeltaVision Elite microscope system built on an Olympus base with a 100 x 1.39 NA and 60 x 1.42 NA objective in an environment chamber heated to 37°C. This system uses a scientific CMOS camera and DV Insight solid state illumination module with the following excitation wavelengths: DAPI = 390/18 nm, FITC = 475/28 nm, TRITC = 542/27 nm, and Alexa 647 = 632/22 nm. Single band pass emission filters had the following wavelengths: DAPI 435/48 nm, FITC = 525/48 nm, TRITC = 597/45 nm, and Alexa 647 = 679/34 nm. A Nikon TI-2 microscope system with 100 x 1.45 NA, 60 x 1.42 NA, and 40 x 0.60 NA objectives in an environment chamber heated to 37°C. This system uses an ORCA-Fusion C14440 digital CMOS camera and Lumencor Spectra light engine with the following excitation wavelengths: DAPI= 390/22 nm, FITC= 475/28 nm, Cy3= 555/28 nm, and Cy5= 637/12 nm. Single band emission filters had the following wavelengths: DAPI = 432/36 nm, FITC = 515/30 nm, Cy3 = 595/31 nm, Cy5 = 680/24 nm. Image acquisition speeds for the live cell imaging were determined on a case-by-case basis and are indicated in the figure legends. For multiple focal planes, a Z - step of 0.2 um (fixed and pulse chase) or single Z - slices (live videos) were used. Images in figures represent a maximum intensity Z - slice projection unless stated otherwise. Brightness and contrast normalized unless noted in figure legend.

### Immunofluorescence assays

Coverslips were fixed in 4% paraformaldehyde (Electron microscopy sciences, Catalog#: 15714) in 1 x PBS for 15 minutes at room temperature. Coverslips were washed 3 x times in PBS before permeabilization with 0.25% Triton-X 100 (ThermoFisher Catalog#: 28314) for 15 minutes. Coverslips were washed 3 x times in PBS and blocked in 2% BSA (Fisher Catalog# BP9703) in PBS for 20 minutes. Antibodies were diluted in PBS according to table S3. Blocking buffer was aspirated and primary then secondary antibodies added for 30 minutes, with 3 x PBS washes between primary and secondary. 10µM DAPI (ThermoFisher Catalog#: D1306) diluted in PBS was added to samples for 10 minutes followed by 3 x PBS washes. Coverslips were mounted onto slides with Prolong Diamond (ThermoFisher Catalog#: P36970) or Vectashield Plus (Fisher Catalog#: H19002), and imaged after drying overnight (Prolong) or immediately (Vectashield Plus).

### Ultrastructure Expansion Microscopy

Ultrastructure expansion microscopy samples were prepared following protocol outlined in Liffner et al. 2023 with the following modifications. Parasites grown in HFFs on 12mm round coverslips (FisherScientific Catalog#: NC1129240) in 6-well plates (Corning Catalog# 3506) were fixed in 4% paraformaldehyde in 1 x PBS for 20 minutes at room temperature. Coverslips were washed 3 x times in PBS and then incubated in 1.4% paraformaldehyde/2% acrylamide (Sigma Catalog#: A4058) in 1 x PBS overnight at 37°C. Coverslips washed in 1 x PBS. To create the expansion gel, parafilm squares were placed in 12-wells on ice and 90 µl monomer solution (19% Sodium Acrylate (Catalog#: 408220), 10% acrylamide, 0.1% N,N’-methylenebisacrylamide (BIS, Sigma Catalog# M1533)) was mixed with 5 µl of 10% tetraethylenediamine (TEMED, ThermoFisher Catalog#: 17919) and 5µl of 10% ammonium persulfate (APS, ThermoFisher Catalog#: 17874), added to parafilm and coverslips pre-dabbed dry were quickly added to gelation solution, cells facing towards drop. Coverslips incubated 5 minutes on ice followed by 1 hour incubation at 37°C, followed by incubation in denaturation buffer (200mM Sodium dodecyl sulphate (SDS), 50mM Tris, 200mM NaCl, in nuclease-free water, pH 9) for 15 minutes at room temperature. Gels were transferred to 1.5ml Eppendorf tubes filled with denaturation buffer and incubated 95°C for 90 minutes. Gels cooled to room temperature before transferring to petri dish with deionized (di) water. Gels were washed three times in diWater with a 30 minutes incubation. The gel diameter was measured to calculate the expansion factor. Gels were placed into 1 x PBS for 30 minutes on room temperature shaker. Gels were cut into quarters and added to 12 wells containing 2% BSA in 1 x PBS. Gels blocked for 30 minutes before incubation with primary antibody (Table S3) diluted in 2% BSA overnight on a shaker at room temperature. Gels washed 4 x times in 0.5% Tween-20 in 1 x PBS (PBS-T), 10 minutes per wash. Secondary antibody (Table S3) and DAPI in 2% BSA added to gels and incubated for 2.5 hours on shaker at room temperature. Gels washed 4 x times in PBS-T, then added to petri dish with deionized water. Two fresh water changes were made 30 minutes apart, then gel cut into square and placed on treated Mattek dish (MatTek dish pre-incubated for 2 hours at 37°C in 0.1mg/ml Poly-D-Lysine (Gibco Cat #: A3890401) and washed 3 x times in water then allowed to air dry). Gels imaged in Mattek dishes immediately using 60 x objective.

#### Live imaging of TgBipA-iKO::APT1_emFP parasites transfected with FNR-RFP plasmid

Control or rapamycin treated parasites were transfected with an FNR-RFP expression plasmid (Table S2), as described above and grown for 24 hours in MatTek dishes containing confluent HFF monolayers. Prior to imaging, dishes were washed three times in 1x PBS and then imaged in fluorobrite DMEM. Imaging speed was 300ms on a single Z-slice.

### TgBipA Co-immunoprecipitation pulldown

Parasites were grown in HFFs in 5 T75 cm^2^ flasks (ThermoFisher Catalog#: 12556010) and freshly egressed parasites were scrapped and centrifuged at 1260 x g for 4 minutes. Pellets were resuspended in 10ml 1 x PBS and counted, then centrifuged again at 1260g for 4 minutes. Pellets were resuspended in 5ml of 1x Lysis buffer (1M imidazole, 1M KCl, 100mM EGTA, 0.5M MgCl_2,_, 0.125mM GTP (Millipore Catalog#: GE27207601), 2mM DTT, 1% TX-100, 0.5 mM PMSF, 10 µg/ml leupeptin, 10 µg/ml pepstatin, 10 µg/ml aprotinin, 10 µg/ml calpeptin, 1.6 mg/ml benzamidine, 50 µl protease inhibitor cocktail (Sigma Catalog#: P8340) in final volume of 11 ml in water) and incubated on ice for 10 minutes. Samples were centrifuged at 17,000 x g for 30 minutes at −4°C and supernatant collected. Supernatant was incubated with 25µl of Anti-FLAG magnetic beads (FisherScientific Catalog#: PIA36797) for 1 hour on a shaker at −4°C, and then magnetic beads were collected from supernatant in an Eppendorf tube using magnetic separation rack (Cell signaling Catalog#: 7017S). Beads were washed 10 x times in 1 x wash buffer (1M imidazole, 1M KCl, 100mM EGTA, 0.5M MgCl_2_, 0.1mM GTP, 2mM DTT in water) and stored at −4°C until analyzed by Mass-spectrometry.

### Image processing, analysis, and statistics

Graphs made in GraphPad Prism as bar charts or superplots (Lord et al., 2020)

#### TgBipA knockout NEAT protein localization counting

To quantify apicoplast morphology upon BipA knockdown, fields of view containing at least 1 parasite vacuole with at least 2 daughters were imaged, scored and APT1 signal categorized based on a ring-like, disordered, or punctate morphology. Cpn60 was categorized based on a single bright puncta, multi-bright puncta, or dim puncta morphologies. N = at least 100 parasites from 3 independent experiments, phenotype occurrence converted to percent of total. Error bars represent standard error of mean. Statistical significance was determined by the means using a two-tailed Student’s Paired t - test.

#### Apicoplast length measurements

To determine if there was a change in apicoplast length during division, apicoplast length was measured on max intensity Z - projections of images with parasites containing an elongated or u-shaped apicoplast using Cpn60 signal and the segmented line tool in Fiji. N = 25 parasites from 3 independent experiments. Error bars represent standard error of mean. Statistical significance was determined by the means using a two-tailed Student’s paired t-test.

#### Expansion apicoplast area measurements

To quantify the reduction in apicoplast size in the TgBipA KO, Z - slices containing non-elongated APT1 signals in an apicoplast ring-shape had their area measured using the polygon selection tool in Fiji. Due to differing expansion factors across samples, areas were adjusted to dataset with final expansion factor of x4.167. N = 20 parasites from 3 independent experiments. Error bars represent standard error of mean. Statistical significance was determined by the means using a two-tailed Student’s paired t-test.

#### Quantification of pulse-chase assay

To quantify the percent of the Jf646 signal which overlaps with the TMR max intensity Z – projections were created. Threshold images were generated using the default thresholding setting in Fiji, then converted to a binary mask and mask area was calculated using analyze particles tool. Percent overlap of Jf646 signal with TMR signal was calculated using the formula:

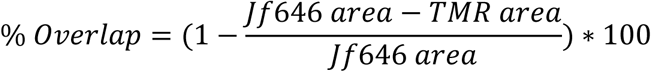

N = 20-30 vacuoles from 3 independent experiments. Error bars represent standard error of mean. Error bars represent standard error of mean. Statistical significance was determined by the means using a two-tailed Student’s Paired t - test.

#### Apicoplast inheritance

To calculate the % of apicoplast being partitioned into dividing daughters, integrated density (iden) measurements using a 25×25 circle were performed for Cpn60 and APT1 on max intensity Z - projections of daughters (daughter A and daughter B) with visible IMC1 staining and either elongated, u-shaped, or newly partitioned APT1 and Cpn60 signal. % of signal inherited by daughter was calculated using:

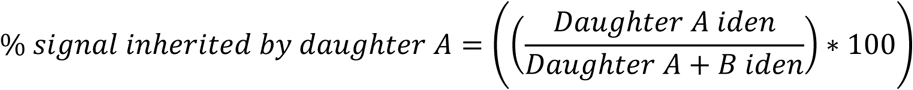

Ratio of inheritance was calculated using:

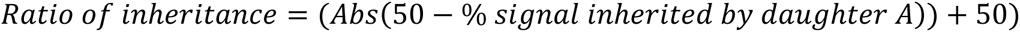

This equation provides a ratio in which the first number is between 50 and 100. N = 20 daughters from 3 independent experiments. Error bars represent standard error of mean. Statistical significance was determined by the means using a two-tailed Student’s paired t-test.

#### Bleach Correction

Time-lapse images and movies were bleached corrected using the histogram matching method in FIJI.

#### Deconvolution

Deconvolution was performed using Softworx v7.2.2 (Deltavision Elite) and the Richardson-lucy method (Nikon Ti-2). All images in figures are deconvolved unless stated otherwise.

#### Expression and Purification of TgBipA from bacteria

An N-terminal his-tagged truncated TgBipA construct was ordered from Twist Biosciences by inserting *E. coli* optimized cDNA encoding residues 518-1333 of TgBipA into a pET28a vector. The Lys536Ala (K536A) variant was constructed using a QuikChange site-directed mutagenesis protocol using primers listed in Table S1. Plasmids were confirmed with Sanger sequencing. Wild-type and K536A mutant recombinant proteins were expressed in *E. coli* BL21(DE3) cells. Cells were grown at 37°C in Luria-Bertani medium supplemented with 30 μg/mL kanamycin to mid-log phase and induced with 0.5mM isopropyl β-D-thiogalactopyranoside (IPTG). The cells were grown an additional 12-15 h at 16°C and harvested by centrifugation (5000 x g for 30 min at 4°C). Cells were washed in 200mM NaCl, 20mM Tris-HCl (pH 7.5) pelleted by centrifugation, and stored at −20°C.

Protein purification was done at 4°C. Cell pellets were resuspended in 20mM HEPES, 20mM imidazole, 400mM KCl, 1mM DTT, 1% glycerol and lysed by sonication, and the lysate clarified by centrifugation (20000 x g for 30 min at 4°C). The supernatant was then loaded onto a HisTrap FF crude column (Cytiva LifeSciences). TgBipA was eluted with an imidazole gradient from 0.02 to 0.4 M in the same buffer. His-tagged protein was concentrated to < 2 mL using an Amicon Centrifugal Filter Unit (Millipore) and applied to a Superdex 200 16/60 prep grade column (Cytiva LifeSciences) equilibrated in 20mM Hepes, 250mM KCl, 1mM EDTA, 1mM DTT, 1% glycerol. Fractions were analyzed by sodium dodecyl sulfate-polyacrylamide gel electrophoresis (SDS-PAGE), and those containing protein that was >95% pure were pooled. Protein quantification was done with the Pierce BCA Protein Assay (ThermoFisher; Catalog #23225).

#### GTP Hydrolysis End Point Assays

The GTPase activity of TgBipA was measured using a malachite green endpoint colorimetric assay^89^. TgBipA (1μM) was incubated with 3mM GTP at 37°C for three hours. To stop the reaction, 30 ml of the reaction mixture was added to 0.8 ml of acidic malachite green solution. After incubation for 30 minutes at room temperature, color formation was assessed at 660 nm using a ThermoFisher Scientific Genesys 150 UV-visible Light Spectrophotometer. Background GTP hydrolysis rates were determined for 3 mM GTP. Reactions were performed at least two times. Experiments were performed in 20mM HEPES, 150mM KCl, 20mM MgCl, 1mM DTT.

## Acknowledgements

We thank Giancarlo Montovano for his assistance in producing recombinant TgBipA protein and members of the Heaslip and Robinson labs for helpful discussions as this work was being carried out. We thank Drs Jeremy Balsbaugh and Jennifer Liddle at UConn Proteomics and Metabolomics for their assistance with the mass spectrometry analysis. Dr. Sabina Absalon (Indiana University) for assistance with the expansion microscopy experiments. Dr. Boris Streiepn (University of Pennsylvania) for providing the anti-Cpn60 antibody. Dr. Christopher de Graffenried (Brown University) for sharing the anti-Ty hybridoma cell line.

