## Supplemental Data File for "An apicoplast localized GTPase is essential for *Toxoplasma gondii* survival"

```
                               G1 Motif
TgBipA       516 RKGGELELRNVAIIAHVDHGKTTLVDALLVHAAELQSASDLRTSWDLEKRKSMQRMMDTG  575
StBipA         1 ---MIENLRNIAIIAHVDHGKTTLVDKLLQQSGTFDAR-----------AETQERVMDSN  47
                       :***:*************** ** ::. :::             :: :*:**:.

PONDR-FIT
                                            G3 Motif
TgBipA       576 QLEKERGITITAKVCSLRFKDGRKINIIDTPGHADFSGEVERVMHLADGVLLVVDAVEGC  635
StBipA        48 DLEKERGITILAKNTAIKWND-YRINIVDTPGHADFGGEVERVMSMVDSVLLVVDAFDGP  105
                 :********* **  :::::*  :***:********.******* :.*.*******.:*

PONDR-FIT
                                      G4 Motif
TgBipA       636 KPQTRVVLQKALQAGLRAVVVVNKVDRNSARPEDVAAQVFDLFLQLDASEEQAEFEVVYA  695
StBipA       106 MPQTRFVTKKAFAHGLKPIVVINKVDRPGARPDWVVDQVFDLFVNLDATDEQLDFPIIYA  165
                  ****.* :**:  **: :**:***** .***: *. ******::***::** :* ::**

PONDR-FIT

TgBipA       696 SALNRQSGMSPSSLVSSMDPLVNAIMRLPSPRERREAHLASLSPSSPSSPSPASPPSPSS  755
StBipA       166 SALNGIAGLDHEDMAEDMTPLYQAIVDHVPA-----------------------------  196
                 ****  :*:. ..:...* ** :**:

PONDR-FIT

TgBipA       756 AGVGGSTPGGLLQMQIAHVDRNAYRGVMAMGKLLGGTLIPGMPVGLQRPGEPLRKATFTG  815
StBipA       197 ---PDVDLDGPLQMQISQLDYNNYVGVIGIGRIKRGKVKPNQQVTIIDSEGKTRNAKVGK  253
                     .   .* *****:::* * * **:.:*::  *.: *.  * :       *:*..

PONDR-FIT

TgBipA       816 VFEYDGMDLRELSITASGGSPTESNVLSASSLSPSSSLSPSSSLSPSSSVSPSSSFSSSS  875
StBipA       254 VLTHLGLER---------------------------------------------------  262
                 *: : *::

PONDR-FIT

TgBipA       876 SVSSSLLASDCSSSLLGGSATEGVAERGPEEEQKEAFPRSENLAVVRSASSLPGDILVLA  935
StBipA       263 ---------------------------------------------IDSNIAEAGDIIAIT  277
                                                              : *  :  ***:.::

PONDR-FIT

TgBipA       936 GVSTAVRVGDSVVDLVDPRPLAPMKVADPSVSIQVLVNTSPLAGKEVETPTSGTMLRQRL  995
StBipA       263 GLG-ELNISDTICDPQNVEALPALSVDEPTVSMFFCVNTSPFCGKEGKFVTSRQILDR-L  335
                 *:.  :.:.*:: *  : . *  :.* :*:**: . *****:.*** :  **  :* : *

PONDR-FIT

TgBipA       996 LAMAERDVSLRVDTNPDEDGMQSAGGCSCLLRLSGRGPLHLALVLETLRREGFELLVAAP 1055
StBipA       336 NKELVHNVALRVEETEDA----------DAFRVSGRGELHLSVLIENMRREGFELAVSRP  385
                      ::*:***: . *             :*:**** ***::::*.:******* *: *

PONDR-FIT
```

```
TgBipA     1056 SVLPRWGDDGELLEPVEEMEVAVPQEHIGTIINEVKQRKGDLLEVVSAASSSASALSTLI 1115
StBipA      386 KVIFRE-IDGRKQEPYENVTLDVEEQHQGSVMQALGERKGDLKNMNPDG----KGRVRLD  440
                .*: *   **.  ** *:: : * ::* *:::: : :***** ::   .    ..   *

PONDR-FIT

TgBipA     1116 FRLPTRKAFGLRSSILTASKGTASIHAASGGYQTVPKGEETDGLVKIARKDRHAIMGSSA 1175
StBipA      441 YVIPSRGLIGFRSEFMTMTSGTGLLYSTFSHYDDIRPGEV--------------------  480
                : :*:*  :*:**.::* :.**. :::: . *: :  **

PONDR-FIT

TgBipA     1176 EFAKRKEKSGSASKGCNAFIPKEGAKQSGGFNIKRGIEGAGGGKKKDRGFLVATEEGTVT 1235
StBipA      481 -------------------------------------------GQRQNGVLISNGQGKAV  497
                                                            :::.*.*::. :*...

PONDR-FIT

TgBipA     1176 GKGALSAQERGQLFVAPGDQVYGGMIVGLNSRGGDLPINVCKAKKLTNMRAAHKEISEGI 1295
StBipA      498 AFALFGLQDRGKLFLGHGAEVYEGQIIGIHSRSNDLTVNCLTGKKLTNMRASGTDEAVIL  557
                . . :. *:**:**:. * :** * *:*::**..** :*  ..********: .: :  :

PONDR-FIT

TgBipA     1196 VPPIEVTLDYGMEIIGANEVMEVTPRSIRLAVRSAMRR------------           1333
StBipA      498 VPPIKMSLEQALEFIDDDELVEVTPTSIRIRKRHLTENDRRRANRGQKEE            607
                ****:::*: .:*:*. :*::**** ***:  *   ..

PONDR-FIT
```

**Figure S1**. Amino acid sequence comparison between TgBipA (TgME49_301380) and *S. enterica* BipA done with the Clustal Omega multiple sequence alignment algorithm[52]. The two proteins are 41% identical with those residues colored red. Residues that are strictly conserved are designated with a star, similar residues with a vertical double dot. The G motifs, defining the G-domain of the protein are highlighted in blue and the conserved loop in the C-terminal domain of the protein, colored green. In addition, the intrinsically disordered residues in TgBipA, predicted using PONDR-FIT[55], are designated by a grey horizontal line below the amino acid sequences.

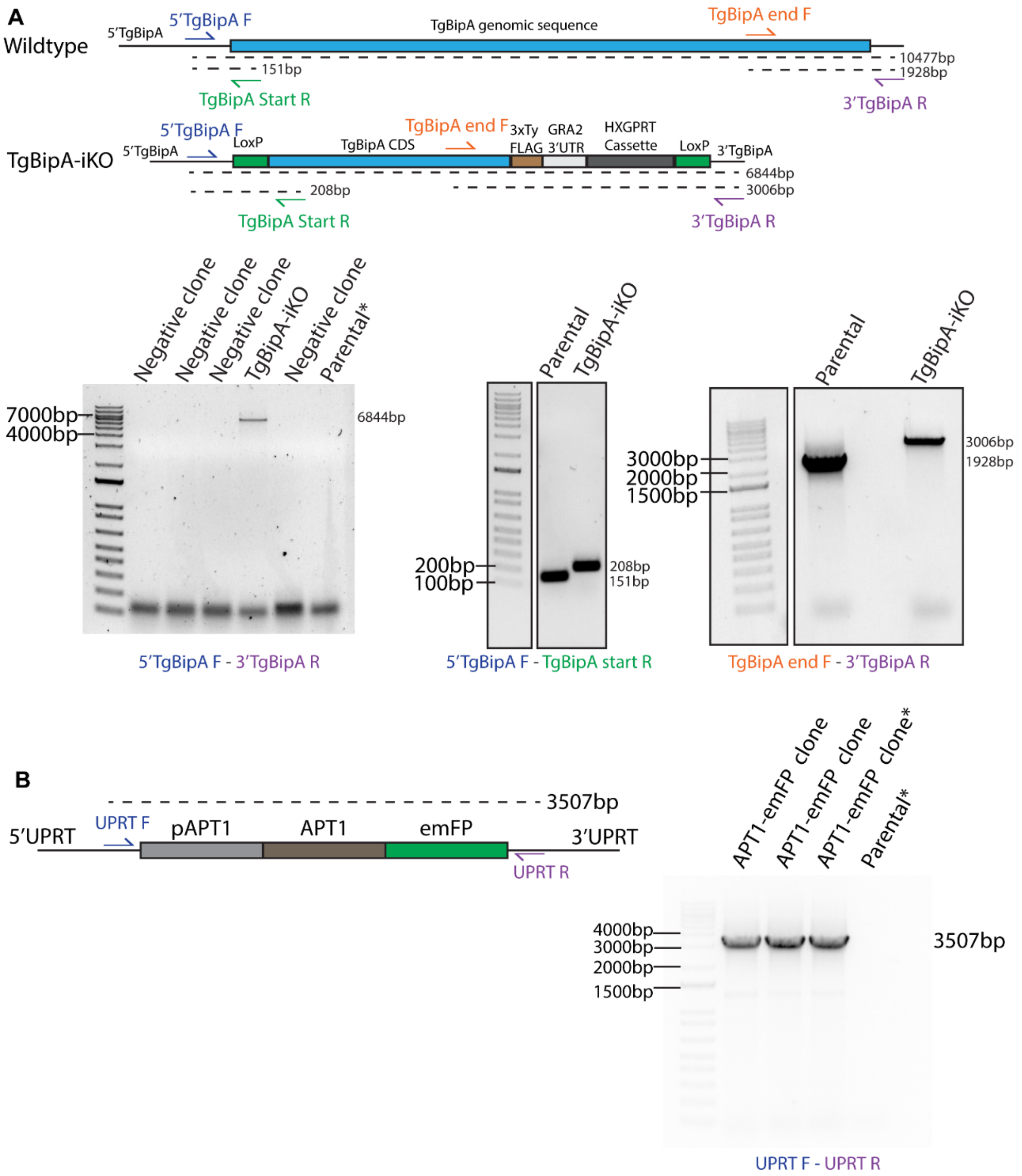


**Figure S2.**

A) *Upper:* Graphic representation of the TgBipA genomic loci in parental and iKO parasite lines respectively. Primer binding sites are indicated. *Lower:* Genomic PCRs for TgBipA-iKO and parental parasite lines. For 5'TgBipA – 3'TgBipA PCR, predicted size of parental loci is 10500bp and yielded no product (Parental* lane). Middle and right gel images are cropped to remove unneeded lanes, cropping denoted by black border. B) Genomic PCR confirmation for insertion of pAPT1-emFP into the UPRT locus of

TgBipA-iKO parasite line clones. Positive clone used in this study denoted by asterisk. Predicted size for wildtype UPRT loci is 5000 base pairs and yielded no product (Parental* lane).

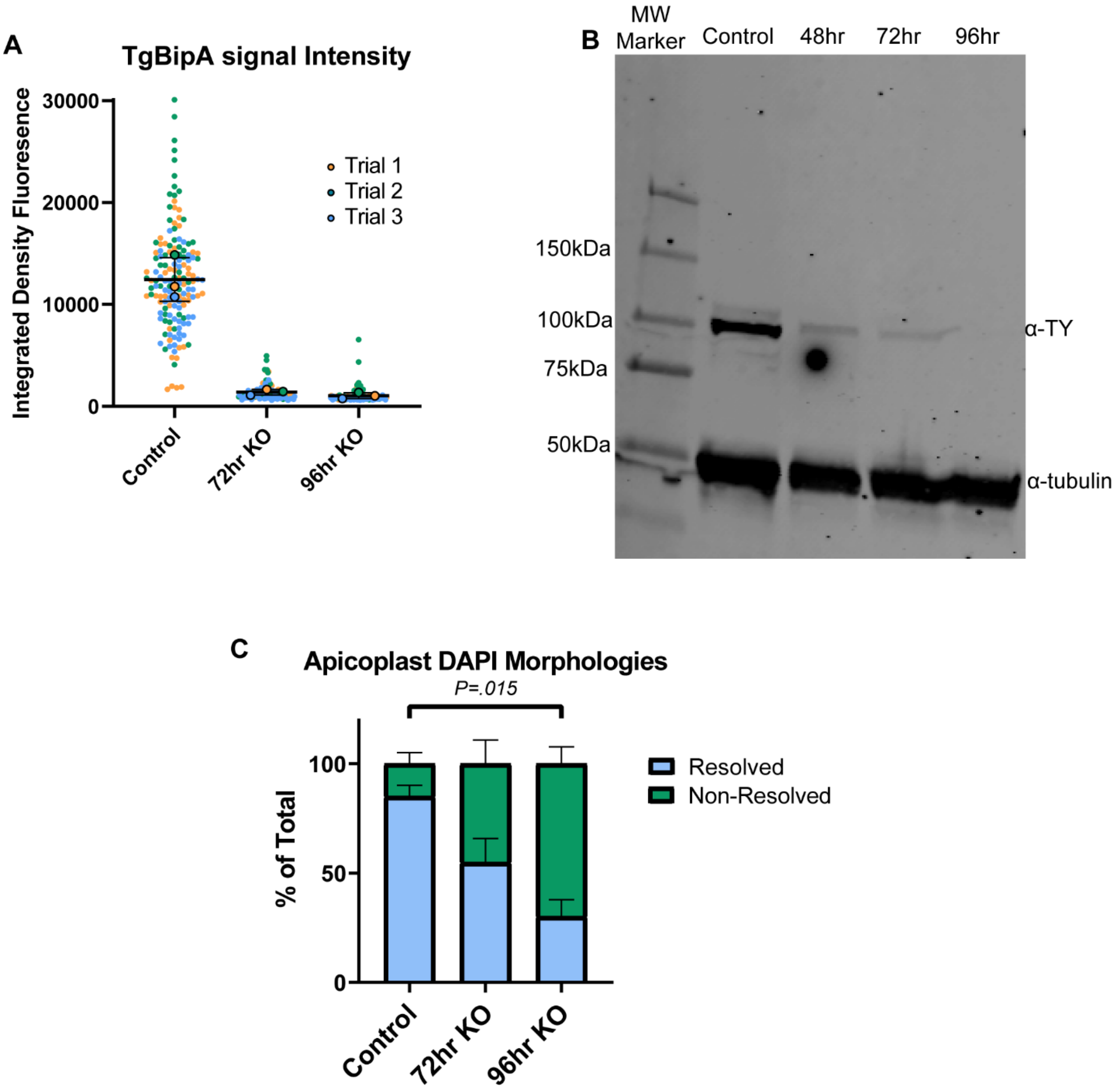


**Figure S3.**

A) TgBipA signal intensity in IF from experiment shown Fig. 2C, n of 50. B) Full western blot from Fig. 2A. C) % of parasites with resolvable apicoplast DAPI signal as determined IF for experiment shown in Fig. 3A and 3B.

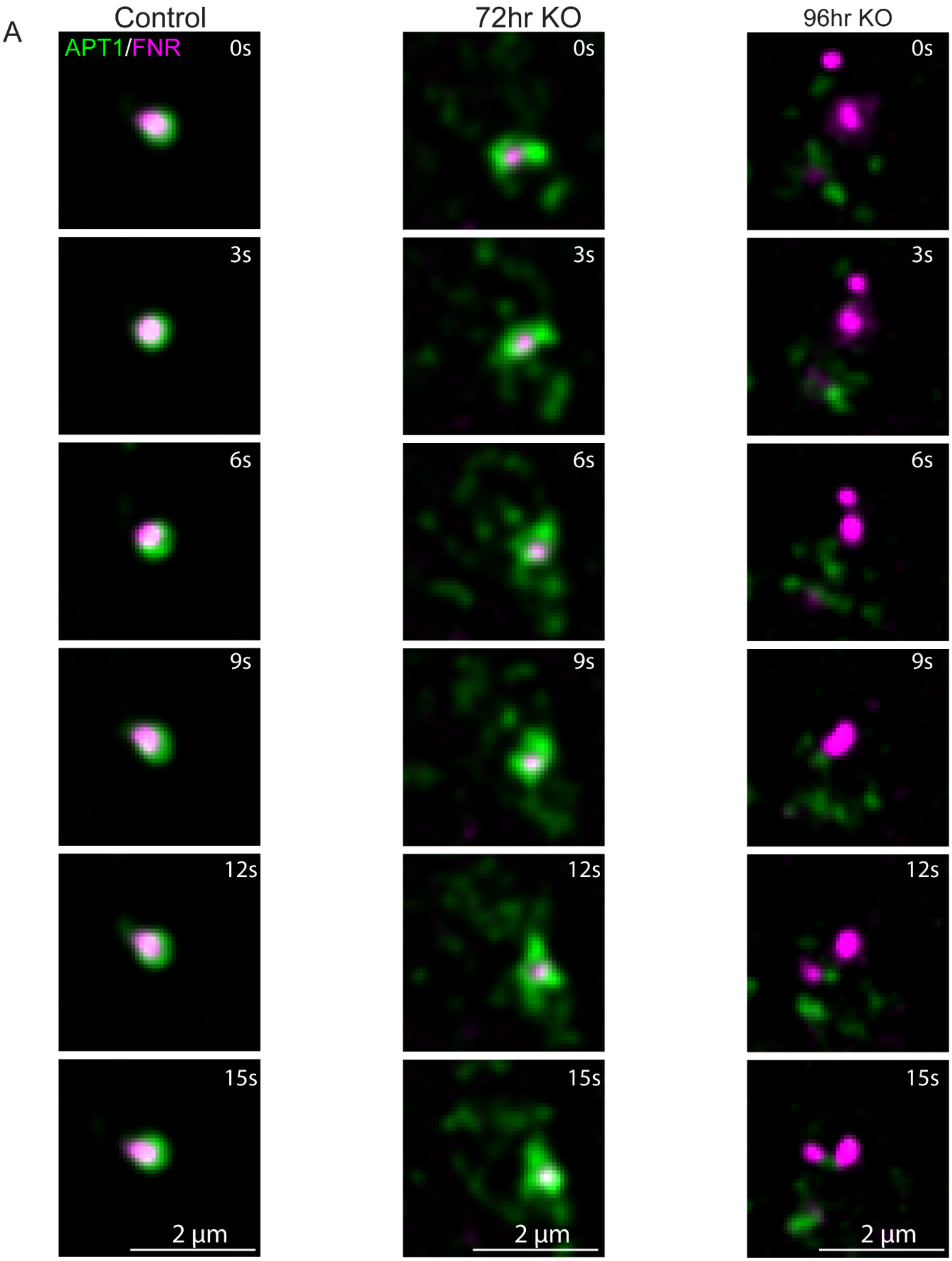

A
Control
72hr KO
96hr KO
APT1/FNR
0s
3s
6s
9s
12s
15s
2 µm

**Figure S4.**

Inset time lapse images from videos 1-3 (Control and TgBipA knockouts (72, and 96 hours after rapa treatment) expressing APT1-emfP and FNR-RFP.) Time in seconds is indicated in top right of image.

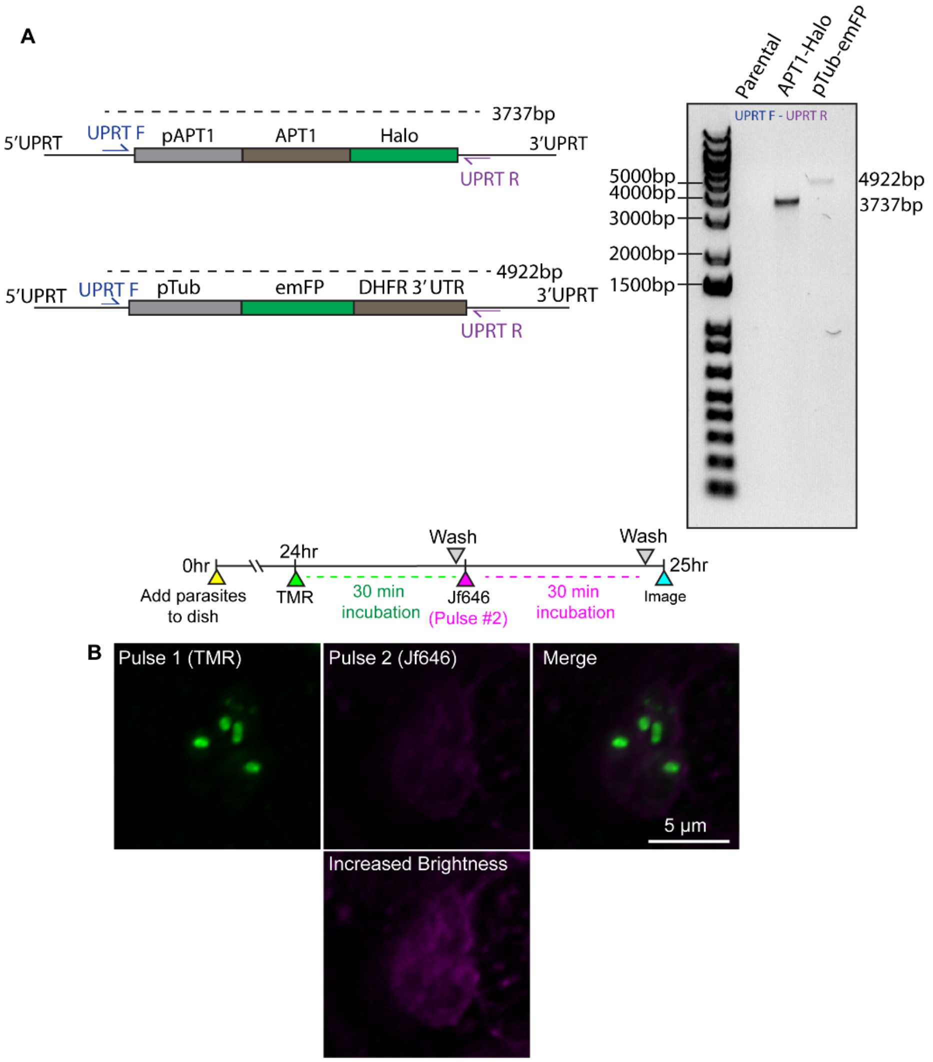


**Figure S5.**

A) Genomic PCR confirmations of pAPT1-Halo and pTub-emFP inserted into the UPRT locus of TgBipA-iKO parasite line. Gel cropped to removed unneeded lanes, denoted by black border. B) Pulse chase assay of TgBipA-iKO::APT1-Halo parasites. Incubation with TMR dye labels Halo protein in the apicoplast. Immediate incubation and imaging with JF646 results in background levels of JF646 staining. Brightness was adjusted in lower panel to show faint Jf646 labeling in the ER. Scale bar = 5 µm.

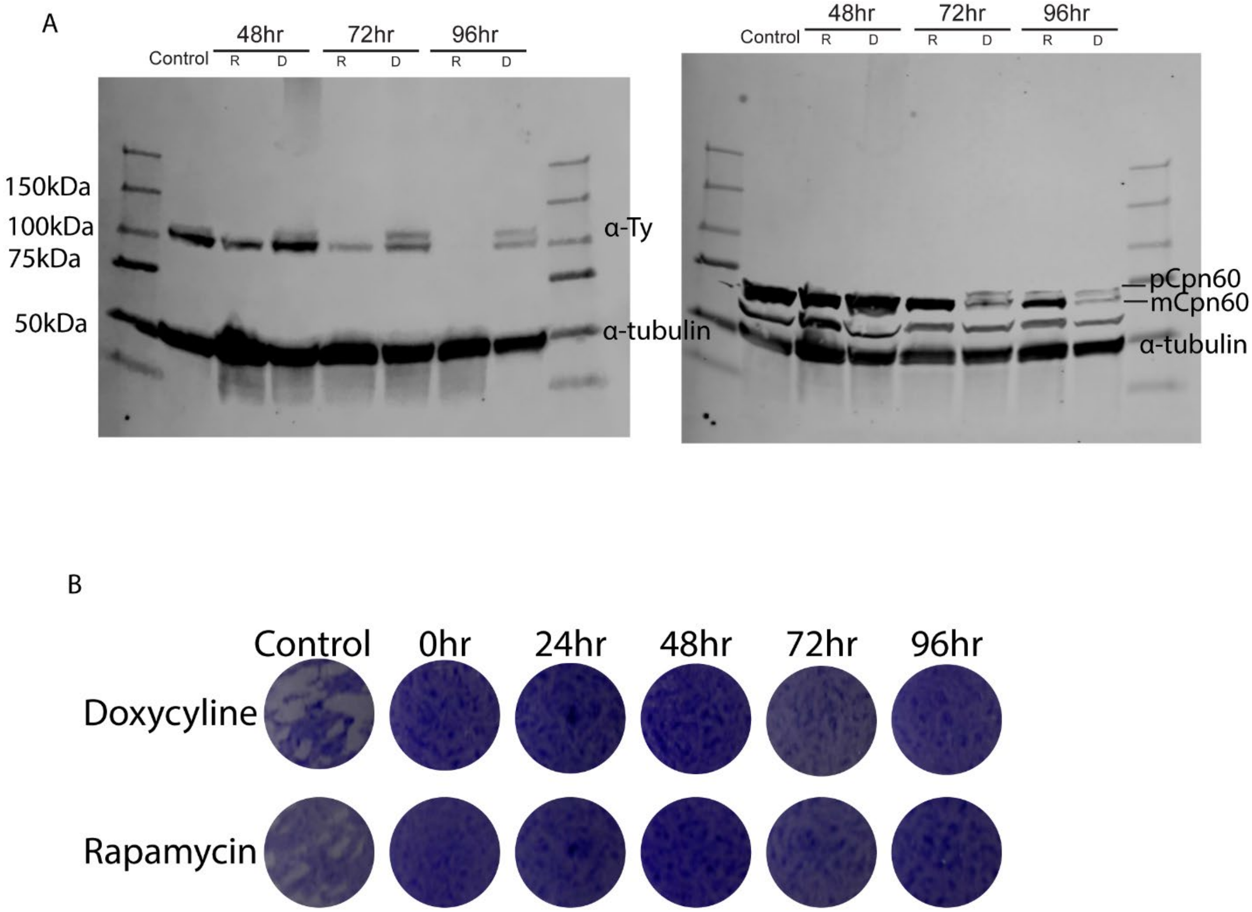


**Figure S6.**

A) Western blots of TgBipA-iKO parasites at 48, 72, and 96 hr post doxycycline or rapamycin treatment. Bands labeled for TgBipA (α-Ty), pre-mature Cpn60 (pCpn60) mature Cpn60 (mCpn60), and Tubulin (α-tubulin). B) Plaque assays of TgBipA-iKO parasites seeded at control, 0, 24, 48, 72, and 96 hr post doxycycline or rapamycin treatment.

A

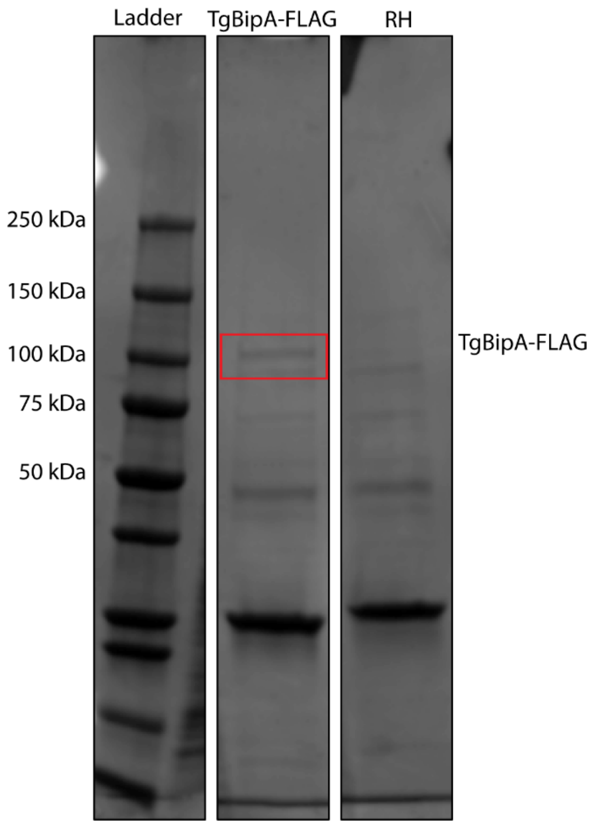


B

**A12345 (100%), 145,367.3 Da**
**Flag-tagged BipA protein OX=508771 GN=RON2 PE=1 SV=2**
**64 exclusive unique peptides, 132 exclusive unique spectra, 331 total spectra, 758/1380 amino acids (55% coverage)**

```
MGDSAVATDV LSLTGGSFSP HTAGEKSNTL SLFLPSSSRP SSRSPPSSAF SPRSSLCVSA GPDCFSLSSG TPASSLCDCF
EEAKEEQESN SRKPKKLCHC QKADAKLRLP SLRPPPPQSP TFLSSSSSSP SFPLSSSSAS SLSFLSALPS SALSSSSVGR
RVKQDLNLKS LFQEHPSSQP RPRHRRTARN FLRVRRRDRS SGTLCKAFSA SRLTLSPRFP RRSFSRCRSV SPAPSGICSL
SVSLYPGPFA KTMASPFPCS SPSSDWSLVS SRFASAPPSR SSAASPHFLP GAFSGASSSS RLSFFSGSNL RQSENCLKRM
RSNSLSENSV SSSLKSLPPS LSPQPISGVA FVAPLSSSSR ASSSLSASLL SPLSYRSASS PRLSVPLLLC LAVLCASVAS
LPLSGSRAPN SRASPLASFF SPCFLSSPLW PSLATSRRAS PLSPSPLSLS PLSASLHLPR GRSRQFLASA PAAALAALRP
AREARAAASS RRPAALFSSA SSVSACPSAE AERRERKGGE LELRNVAIIA HVDHGKTTLV DALLVHAAEL QSASDLRTSW
DLEKRKSMQR MMDTGQLEKE RGITITAKVC SLRFKDGRKI NIIDTPGHAD FSGEVERVMH LADGVLLVVD AVEGCKPQTR
VVLQKALQAG LRAVVVVNKV DRNSARPEDV AAQVFDLFLQ LDASEEQAEF EVVYASALNR QSGMSPSSLV SSMDPLVNAI
MRLPSPRERR EAHLASLSPS SPSSPSPASP PSPSSAGVGG STPGGLLQMQ IAHVDRNAYR GVMAMGKLLG GTLIPGMPVG
LQRPGEPLRK ATFTGVFEYD GMDLRELSIT ASGGSPTESN VLSASSLSPS SSLSPSSSLS PSSSVSPSSS YSSSSSVSSS
LLASDCSSSL LGGSATEGVA ERGPEEEQKE AFPRSENLAV VRSASSLPGD ILVLAGVSTA VRVGDSVVDL VDPRPLAPMK
VADPSVSIQV LVNTSPLAGK EVETPTSGTM LRQRLLAMAE RDVSLRVDTN PDEDGMQSAG GCSCLLRLSG RGPLHLALVL
ETLRREGFEL LVAAPSVLPR WGDDGELLEP VEEMEVAVPQ EHIGTIINEV KQRKGDLLEV VSAASSSASA LSTLIFRLPT
RKAFGLRSSI LTASKGTASI HAASGGYQTV PKGEETDGLV KIARKDRHAI MGSSAEFAKR KEKSGSASKG CNAFIPKEGA
KQSGGFNIKR GIEGAGGGKK KDRGFLVATE EGTVTGKGAL SAQERGQLFV APGDQVYGGM IVGLNSRGGD LPINVCKAKK
LTNMRAAHKE ISEGIVPPIE VTLDYGMEII GANEVMEVTP RSIRLAVRSA MRRLIKDYKD DDDKRSSMEV HTNQDPLDGS
EVHTNQDPLD EVHTNQDPLD
```

**Figure S7.**

A) Proteins purified by Flag immunoprecipitation from TgBipA-TyFLAG and RH control lysates were visualized on an SDS-PAGE gel stained with Coomassie. B) TgBipA peptides identified by mass-spectrometry analysis. Identified residues highlighted in yellow, modified residues in green. No peptides were found in the N-terminus of the protein suggesting proteolytic processing.

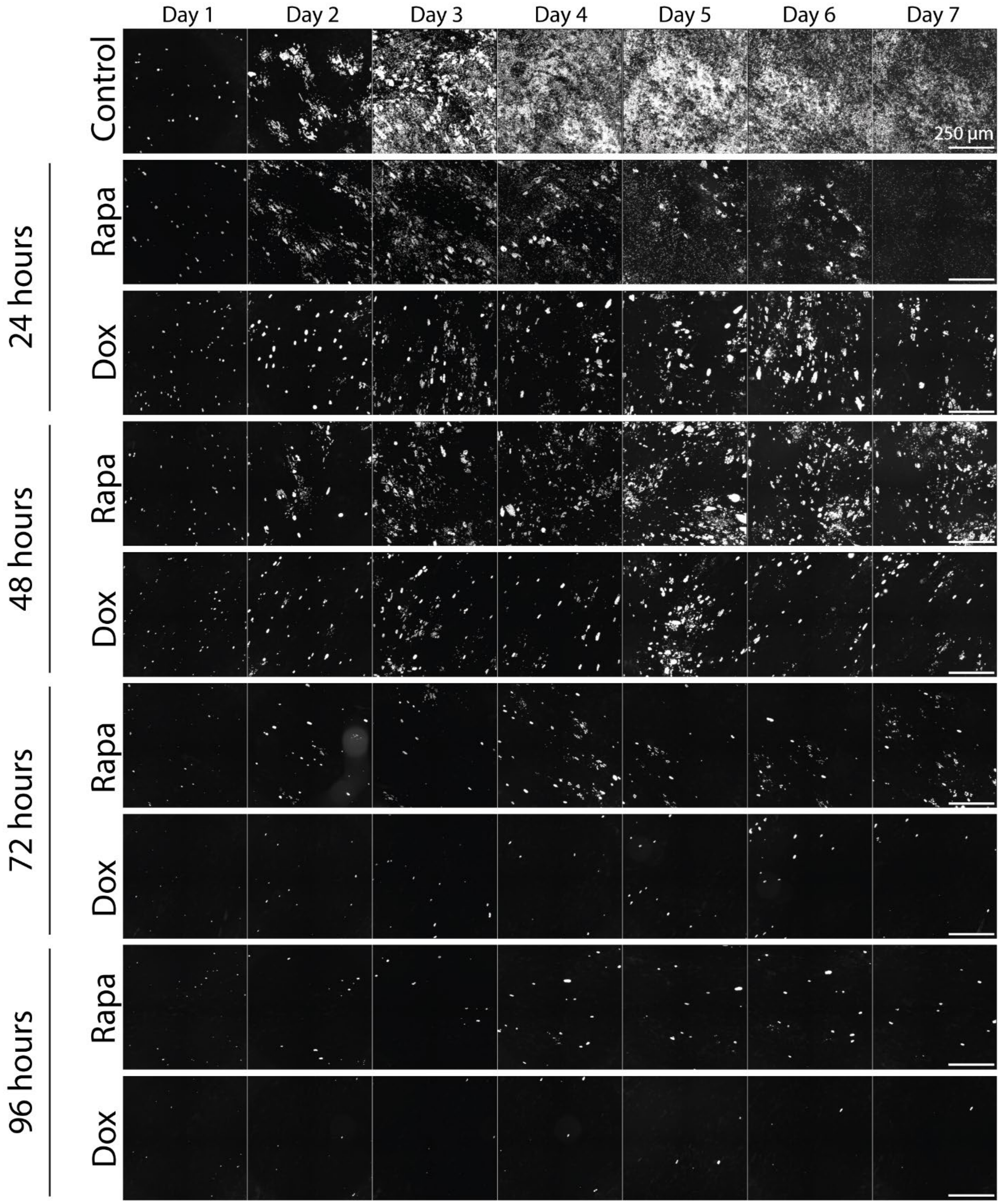
Day 1
Day 2
Day 3
Day 4
Day 5
Day 6
Day 7
Control
250 µm
24 hours
Rapa
Dox
48 hours
Rapa
Dox
72 hours
Rapa
Dox
96 hours
Rapa
Dox

**Figure S8.**

Fluorescence microscopy images of taken every 24 hours during the fluorescence growth assay performed in Figure 5C. Scale bar = 250 μm.

A

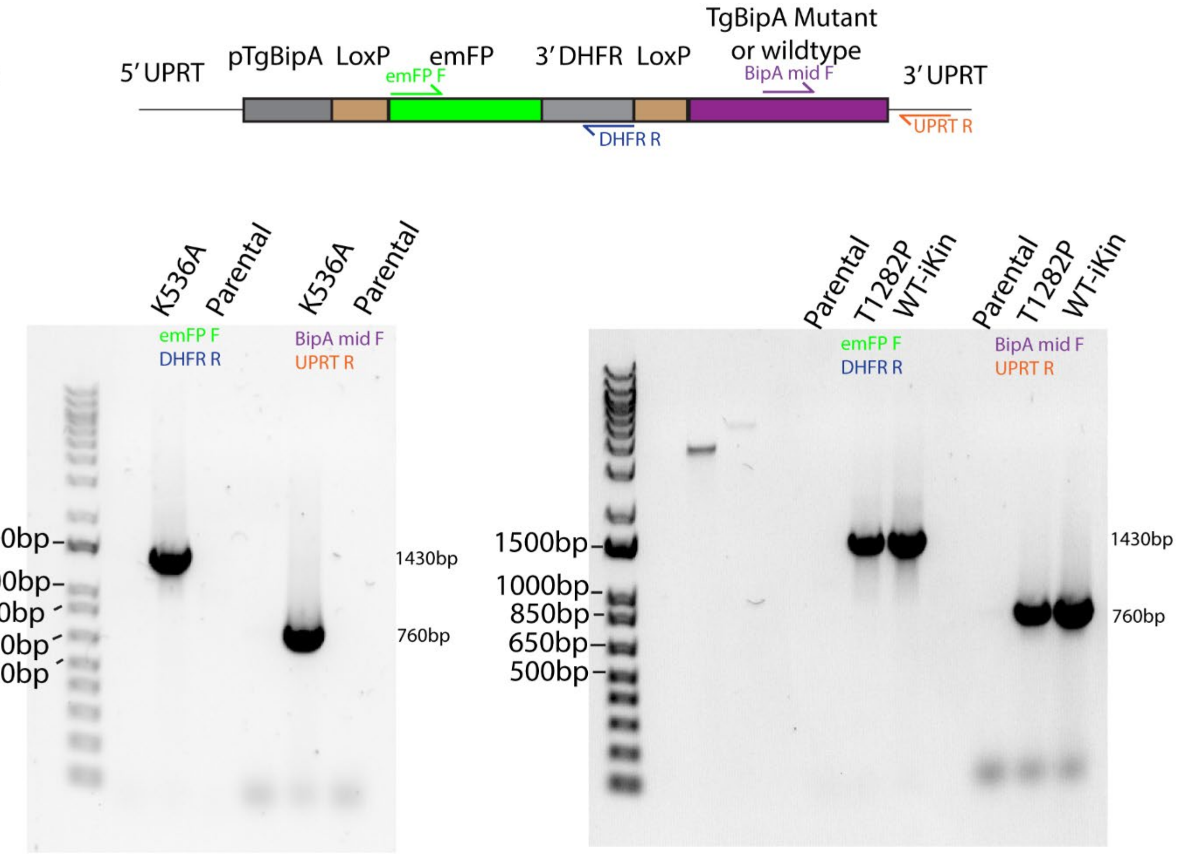
5′ UPRT
pTgBipA
LoxP
emFP
3′ DHFR
LoxP
TgBipA Mutant or wildtype
3′ UPRT
emFP F
BipA mid F
DHFR R
UPRT R
K536A
Parental
K536A
Parental
emFP F
DHFR R
BipA mid F
UPRT R
1500bp
1000bp
850bp
650bp
500bp
1430bp
760bp
Parental
T1282P
WT-iKin
Parental
T1282P
WT-iKin
emFP F
DHFR R
BipA mid F
UPRT R
1500bp
1000bp
850bp
650bp
500bp
1430bp
760bp

B

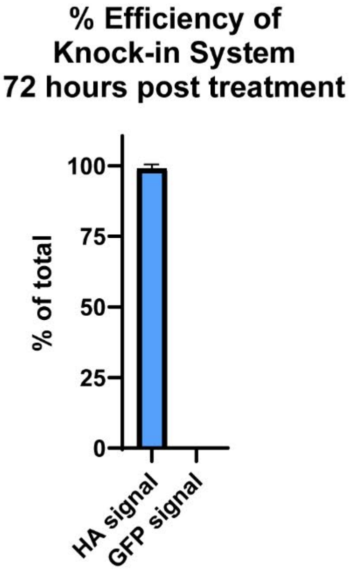
% Efficiency of Knock-in System 72 hours post treatment
100
75
50
25
0
% of total
HA signal
GFP signal

C

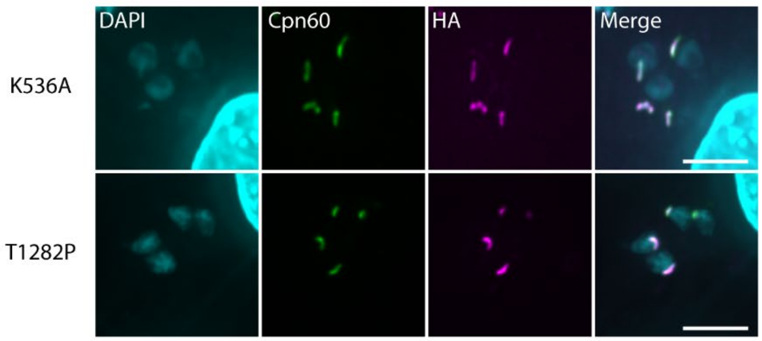
DAPI
Cpn60
HA
Merge
K536A
T1282P

**Figure S9:**

A) Genomic PCR confirmations of TgBipA-iKin mutant rescue lines. B) % of parasites expressing HA-tagged construct which have lost GFP signal 72 hours after rapamycin treatment. n of 100 parasites in 2 independent experiments. C) Immunofluorescence images of K536A-ikin and T1282P-ikin parasites 24 hours after rapamycin treatment show mutations do not alter apicoplast localization. Scale bar = 5 μm.

**Video S1.**

Control TgBipA-iKO parasites expressing APT1-emFP and FNR-RFP imaged live for 30 seconds. Imaging speed 3 frames/sec. playback is 2x realtime. White box denotes inset region in Fig. S4. Scale bar = 5 μm.

**Video S2.**

ΔTgBipA (72H after rapa treatment) parasites expressing APT1-emFP and FNR-RFP imaged live for 30 seconds. Imaging speed 3 frames/second. Playback is 2x realtime. Brightness is adjusted to allow visualization of APT1 signal. White box denotes inset region in Fig. S4. Scale bar = 5 μm.

**Video S3.**

ΔTgBipA (96H after rapa treatment) parasites expressing APT1-emFP and FNR-RFP imaged live for 30 seconds. Imaging speed 3 frames/second. Playback is 2x realtime. Brightness is adjusted to allow visualization of faint APT1 and FNR signal. White box denotes inset region in Fig. S4. Scale bar = 5 μm.

**Table S1**. List of primers used in this study

**Table S2.** List of plasmids used in this study

**Table S3.** List of antibodies used in this study

**Table S4.** Mass-spectrometry data from anti-Flag pulldowns for TgBipA-FLAG and RH parasite lines. Combined data from 3 independent experiments.
